# The mutational landscape of *Bacillus subtilis* conditional hypermutators shows how proofreading skews DNA polymerase error rates

**DOI:** 10.1101/2023.12.29.573609

**Authors:** Ira Tanneur, Etienne Dervyn, Cyprien Guérin, Guillaume Kon Kam King, Matthieu Jules, Pierre Nicolas

## Abstract

Polymerase errors during DNA replication are a major source of point mutations in genomes. The resulting rate of spontaneous mutation also depends on the counteracting activity of DNA repair mechanisms, with mutator phenotypes appearing constantly and allowing for periods of rapid evolution in nature and in the laboratory. Here, we use the Gram-positive model bacterium *Bacillus subtilis* to disentangle the contributions of DNA polymerase initial nucleotide selectivity, DNA polymerase proofreading, and mismatch repair (MMR) to the mutation rate. To achieve this, we constructed several conditional hypermutators with a proofreading-deficient allele of *polC* and/or a deficient allele of *mutL* and performed mutation accumulation experiments. With their wide range of mutation rates and contrasting mutation profiles, these conditional hypermutators enrich the *B. subtilis* synthetic biology toolbox for directed evolution. Using mathematical models, we investigated how to interpret the apparent probabilities with which errors escape MMR and proofreading, highlighting the difficulties of working with counts that aggregate potentially heterogeneous mutations and with unknowns about the pathways leading to mutations in the wild-type. Aware of these difficulties, the analysis shows that proofreading prevents partial saturation of the MMR in *B. subtilis* and that an inherent drawback of proofreading is to skew the net polymerase error rates by amplifying intrinsic biases in nucleotide selectivity.

## Introduction

Substitutions, insertions and deletions of single base pairs in genomes can have diverse consequences on encoded molecular functions, from no effect to abrupt change, most often in the direction of deterioration (Eyre-Walker and Keightley 2007). As such, point mutations are a constant threat to the integrity of the genetic information, even if they are also essential to adaptive evolution. In the long term, point mutation rates themselves result from an evolutionary process. An hypothesis, supported by the comparison of mutation rates across the tree of life, postulates that they are simply maintained as low as possible; the limit being the “drift-barrier”, where the strength of selection is matched by the opposite pressure of random genetic drift and mutation, presumably biased towards creating weak mutators (Lynch et al. 2016). Mutation rate might thus have nothing to do with the benefit of evolvability for long-term survival, the energetic cost of fidelity, or a biophysical limit. In bacteria, this rate is already typically as low as one mutation introduced in the genome per one thousand generations, but mutation rates one order of magnitude below those observed in nature were reported to arise under scenarios of artificial evolution (Deatherage et al. 2018; Dervyn et al. 2023).

However, mutator phenotypes occur everywhere and their contribution to evolution is difficult to estimate (Taddei et al. 1997; Couce et al. 2017). In asexual populations, where mutator alleles and the mutations that they generate remain linked across successive generations, mutators are advantageous when the potential for fitness improvement is high. For example, in pathogenic bacteria, mutators have been associated with complex antibiotic resistance (Dulanto Chiang et al. 2022), rapid evolution within the host during infection (Oliver and Mena 2010), and atypical virulence traits (Rudenko et al. 2020). Mutators also emerge spontaneously during laboratory evolution in response to applied selective pressures (Sniegowski et al. 1997; Swings et al. 2017). To foster adaptation, conditional mutator circuits and systems have been engineered for some organisms as part of the synthetic biology toolbox (Badran and Liu 2015; Sherer and Kuhlman 2020; Molina et al. 2022). Understanding the molecular factors that determine mutation rates is therefore of strong fundamental and applied interest.

The sources of spontaneous mutations in living cells are diverse, including DNA lesions caused by endogenous and exogenous agents, errors introduced by the DNA polymerase during replication, and the activity of error-prone polymerases recruited in response to stress (Maki 2002). The resulting mutation rate depends on the intensity of these sources and of the counteracting activity of DNA repair mechanisms, which work in concert to ensure that the correct genetic material is passed on to daughter cells. Two essential mechanisms, conserved from prokaryotes to eukaryotes, ensure accurate repair of both bulky and non-bulky lesions resulting from DNA damage, including those induced by reactive oxygen species, a major source of DNA errors (Foster et al. 2015). NER (Nucleotide Excision Repair) repair various drug- and UV-induced lesions (*i.e.* bulky lesions), while BER (Base Excision Repair) repair lesions caused by various chemical assaults, such as alkylation, oxidation, deamination, etc. (*i.e.* non-bulky lesions).The accuracy of DNA replication depends on three critical mechanisms: the initial selectivity of the DNA polymerase, which is responsible for inserting the correct nucleotide; the proofreading, which removes misincorporated nucleotides through polymerase-associated exonucleases; and the mismatch repair (MMR), which adds a second layer of error correction shortly after replication (Ganai and Johansson 2016).

In bacteria, genome replication is carried out by a multiprotein machine classified into the C-family of DNA polymerase holoenzymes, in which the catalytic polymerase α-subunit exists in two primary forms, DnaE and PolC (Timinskas et al. 2014). A representative example of DnaE is found in the extensively studied Gram-negative model bacterium *Escherichia coli*. Conversely, PolC is predominant in low-GC Gram-positive bacteria, such as *Bacillus subtilis* (Sanjanwala and Ganesan 1991). In this organism, replication elongation involves two essential polymerases, PolC and DnaE (Dervyn et al. 2001), which have distinct functions: PolC does most of the DNA synthesis, but only DnaE can elongate from RNA primers on the lagging strand before passing the DNA fragment to PolC. In the absence of proofreading and MMR, the error rate of the *E. coli* α-subunit replication machinery has been estimated to be approximately 10^-6^ per base pair per generation both *in vitro* (Fujii et al. 1999) and *in vivo* (Niccum et al. 2018). Given this error rate and the size of the *E. coli* genome, about 5 mutations are expected to be introduced per generation.

The exonuclease domain essential for proofreading is encoded as an integral part of the vast majority of PolC polymerases (Timinskas et al. 2014), including *B. subtilis* PolC. In contrast, DnaE polymerases do not have their own exonuclease domain. The proofreading activity of the *E. coli* DNA PolIII holoenzyme containing DnaE is based on an exonuclease domain located in the ε-subunit. Errors made by polymerases devoid of proofreading can also sometimes be corrected by a process known as proofreading in *trans*, or extrinsic proofreading, which is well described between eukaryotic DNA polymerases (Zhou et al. 2021). In *B. subtilis*, data suggest that the PolC exonuclease is able to correct errors made by the error-prone DnaE polymerase (Bruck et al. 2003; Paschalis et al. 2017).

The MMR is a universal mechanism that is responsible for correcting errors that occur during DNA replication and escape proofreading. Upon detection of a replication error, the mismatch-sensing protein, MutS, recruits MutL. Most prokaryotic and eukaryotic MutL homologs, from humans to bacteria, possess a highly conserved endonuclease active site that serves to remove mismatches (Pillon et al. 2010; Bolz et al. 2012). *E. coli* has been the primary model for studying MMR, but its MutL lacks the endonuclease activity encoded here in a distinct protein, MutH, which specifically nicks the unmethylated and thus nascent strand (Lenhart et al. 2012). In the absence of MutH and Dam methylation, the process that guides MutL to the nascent strand remains unclear in most prokaryotes and all eukaryotes (Kadyrov et al. 2006). In bacteria, MMR increases the fidelity of the chromosomal DNA replication pathway approximately 100-fold, and MMR is considered to be for repairing the most frequent replication errors (Lujan et al. 2012). Mutator phenotypes found in nature are often caused by mutations that inactivate the MMR.

Studying organisms such as *B. subtilis*, with a PolC polymerase and an MMR pathway more widely conserved across biology can provide important insights into the coordinated functioning of these systems within living cells (Klocko et al. 2011). Extending previous work characterizing the mutation profiles of MMR-deficient *B. subtilis* strains (Sung et al. 2015; Schroeder et al. 2016), the main goal of our study was to jointly assess the respective contributions of initial nucleotide selectivity, proofreading and MMR to the mutation rates and their interdependencies. For this purpose, we constructed and analyzed several conditional hypermutators. Analysis of the data in light of several mathematical models suggests that proofreading is needed to avoid partial saturation of the MMR, as previously reported in *E. coli* (Schaaper 1988; Niccum et al. 2018), but also that an inherent effect of proofreading is to skew the net polymerase error rate. With their wide range of mutation rates and contrasting mutation profiles, the conditional hypermutators also enrich the *B. subtilis* synthetic biology toolbox for directed evolution.

## Results

### The mutation rate of *Bacillus subtilis* can be increased up to 6,000 times

We have constructed five mutant strains with expected hypermutator phenotypes from a strain derived from *B. subtilis* 168. The first two strains are constitutively MMR deficient as a result of single deletions of *mutS* and *mutL*. The other three were engineered for conditional inactivation of either or both of MMR or proofreading and were therefore expected to be inducible hypermutators (**Figures S1** and **S2**). In these three strains, the IPTG-inducible promoter P*hs* controls the expression of mutant alleles selected for their ability to competitively displace their functional counterparts. The first allele, designated *mutL\**, has a mutation in the region encoding the ATP hydrolysis active site of MutL which has been reported to have a dominant negative effect (Bolz et al. 2012). The second allele, designated *polC\**, encodes a proofreading deficient variant of PolC with a mutation in its exonuclease domain (Sanjanwala and Ganesan 1991). The final strain, which was expected to have the highest mutation rate under full induction, expresses these two deficient alleles in a synthetic operon (*mutL* polC\**). The following notations are used for the reference parental strain derived from 168 and the five mutant strains: R^168^, Δ*L*, Δ*S*, *L*\*, *C*\*, *LC*\*.

Fluctuation assays were performed (**Figure S3**) to compare the rate of mutation to rifampicin resistance of these strains. Point estimates and confidence intervals are shown in **Figure 1**, and results are detailed in **Table S1**. In the absence of IPTG, the estimated mutation rate of R^168^ was 9.74×10^-10^ per generation. Constitutive inactivation of the MMR in Δ*L* and Δ*S* increased the mutation rate by a factor of approximately 85, with no statistically significant difference between the two strains. This is close to the factor of about 60 previously obtained for a double deletion of *mutL* and *mutS* in the *B. subtilis* PY79 genetic background with a fluctuation assay also based on rifampicin (Schroeder et al. 2016). Mutation rates in the absence of IPTG were slightly higher for the inducible strains (*L*\*, *C*\*, *LC*\*) than for R^168^ (up to 1.09×10^-8^ for *LC*\*), likely reflecting the low basal activity already described for P*_hs_* (Guiziou et al. 2016). From there, the mutation rate of the three inducible strains increased with IPTG concentration, reaching a plateau between 50 and 100 µM. At full induction (100 µM IPTG), the mutation rates ranging from 3.66×10^-7^ for *L*\* to 5.78×10^-6^ for *LC*\*. The mutation rate in *L*\* is comparable to or slightly higher than in Δ*L* and Δ*S*, while the mutation rate in *LC*\* represents an approximately 6,000-fold increase over R^168^.

**Figure 1.**
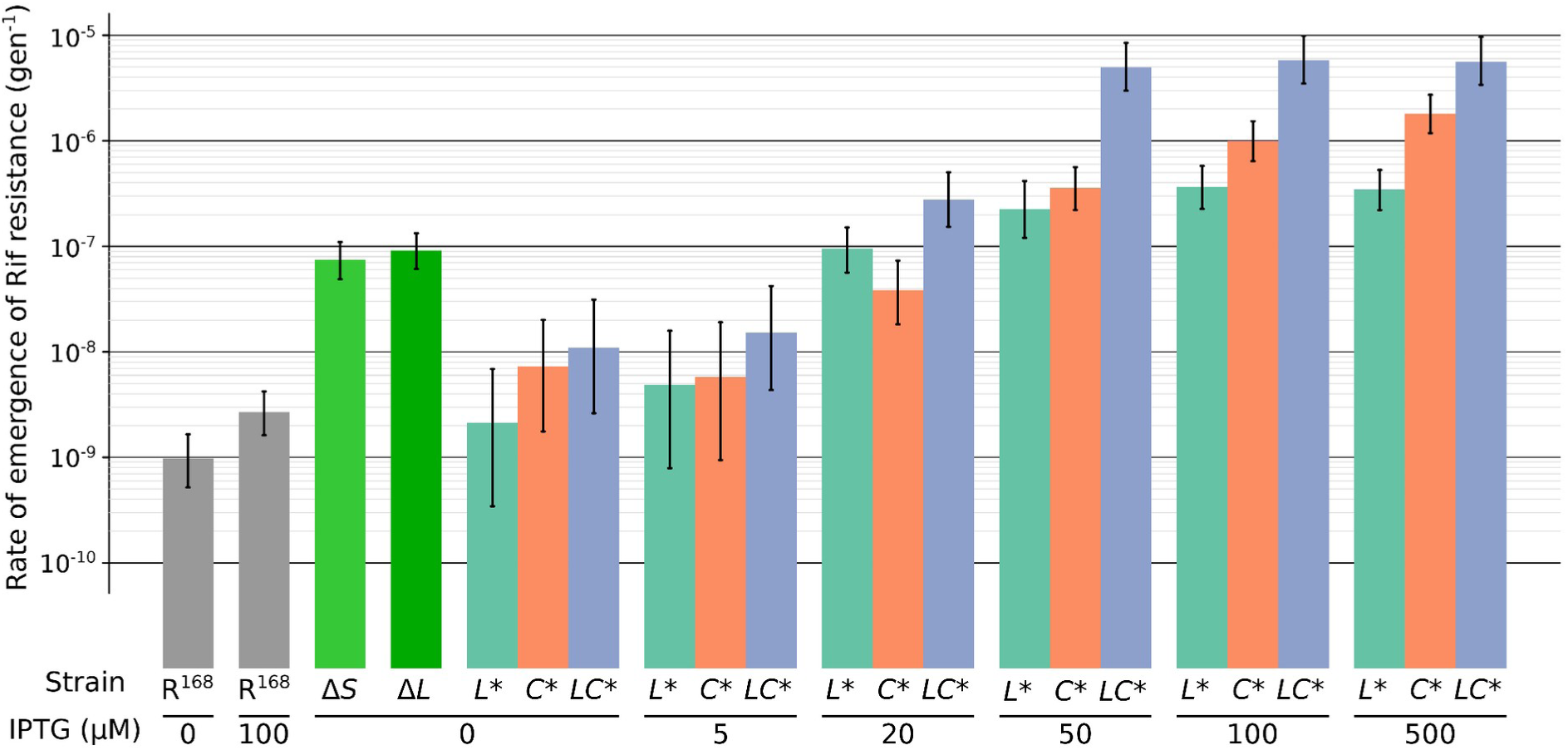
Mutation rate to rifampicin resistance measured by fluctuation assays for increasing IPTG concentration. Each color corresponds to a strain; vertical bars represent 95% confidence intervals.

Much higher mutation rates than in the reference strain may induce stress responses, potentially altering the physiology of each mutant differently. However, we did not detect any substantial impact on growth in 96-well microtiter plates (**Figure S4**). We also performed transcriptomics experiments on the R^168^, *L*\*, *C\** and *LC*\* strains in the presence of 100 µM IPTG. The analyses did not reveal any significantly differentially expressed genes between the strains. However, they did allow us to quantify the expression of the mutant alleles relative to wild-type alleles for *mutL* and *polC*. Upon induction, the mutant allele accounted for 96-98% of the total mRNA pool for the gene in question (**Table S2**), which is consistent with the results of fluctuation assays that give a mutation rate in *L\** as high as in Δ*L* and Δ*S* (*i.e.* complete inactivation of the MMR). We therefore concluded that the mutation profiles of these strains can be attributed to the sole inactivation of the two targeted DNA repair pathways.

### Highest mutation rates are counter-selected in mutation-accumulation experiments

Mutation-accumulation experiments give access to the molecular nature of mutations (Lynch et al. 2016), in contrast to fluctuation assays, which only provide a mutation rate aggregated across a range of mutations that confer a screenable phenotype. For each of the six strains (R^168^, Δ*L*, Δ*S*, *L* *, *C*\*, *LC*\*), four independent mutation-accumulation lines (MA-lines) were propagated by repeated cycles (MA-steps, **Figure S5**) of colony picking, dilution, and plating on LB, in the presence of 100 µM IPTG when relevant. By randomly selecting a single colony, each MA step creates a bottleneck in the propagated population, the whose purpose of which is to limit genetic diversity, thereby maximizing random genetic drift and minimizing natural selection. The interval between two bottlenecks, one MA-step, was estimated to be 25.6 generations on average. We performed whole-genome sequencing at the endpoint of each line; we also sequenced intermediate time-points (up to 4 for *LC*\*) to detect changes in mutation rates (**Figure S5**). A total of 56 clones isolated after 1 to 37 MA-steps were sequenced. Point mutations - substitutions, insertions (ins), and deletions (del) - were identified. Mutation rates per base pair (bp) and generation (abbreviated bp^-1^.gen^-1^) were estimated for each time interval of each MA-line. Previously collected data from mutation-accumulation experiments using *B. subtilis* 3610 (R^3610^) and its corresponding Δ*mutS* mutant strain (Δ*S*^3610^) were also incorporated to increase statistical power (Sung et al. 2015; Sung et al. 2016). The detailed list of all mutations found is provided in **Table S3**.

In the four independent lines of R^168^, only one nucleotide substitution was identified after 37 MA-steps, giving an estimated substitution rate of 7×10^-11^ bp^-1^.gen^-1^ (**Table 1**). This rate was not statistically significantly different from a previous report on *B. subtilis* (Sung et al. 2016), recalculated to 3.4×10^-10^ bp^-1^.gen^-1^ (**Table 1**).

**Table 1.** Aggregated number of substitutions, substitution rate, and proportion of transversions for each strain.

| Strain<sup>a</sup> | Li-<br>nes | Generations<br><sub>b,c</sub> | Substitutions<sup>c</sup> | | | Substitution rate<br>($\text{bp}^{-1}.\text{gen}^{-1}$)<br>[95% CI] | Proportion of<br>transversions<br>[95% CI] |
| --- | --- | --- | --- | --- | --- | --- | --- |
|  |  |  | Total | ts<sup>d</sup> | tv<sup>e</sup> |  |  |
| $R^{168}$ | 4 | 3,790 | 1 | 1 | 0 | $7.0 \times 10^{-11}$ [0.18-39$\times 10^{-10}$] | 0.00 [0.00-0.98] |
| $R^{3610}$ | 49 | 248,920 | 319 | 238 | 81 | $3.4 \times 10^{-10}$ [3.0-3.8$\times 10^{-10}$] | 0.25 [0.21-0.31] |
| $\Delta S^{3610}$ | 19 | 38,000 | 4,844 | 4,711 | 133 | $3.4 \times 10^{-8}$ [3.3-3.5$\times 10^{-8}$] | 0.03 [0.02-0.03] |
| $\Delta L$ | 4 | 2,151 | 149 | 147 | 2 | $1.8 \times 10^{-8}$ [1.5-2.1$\times 10^{-8}$] | 0.01 [0.00-0.05] |
| $\Delta S$ | 4 | 2,151 | 157 | 155 | 2 | $1.9 \times 10^{-8}$ [1.6-2.2$\times 10^{-8}$] | 0.01 [0.00-0.05] |
| *L\** | 4 | 2,151 | 113 | 111 | 2 | $1.4 \times 10^{-8}$ [1.1-1.7$\times 10^{-8}$] | 0.02 [0.00-0.06] |
| MMR-<sup>168</sup> | 12 | 6,453 | 419 | 413 | 6 | $1.7 \times 10^{-8}$ [1.6-1.9$\times 10^{-8}$] | 0.01 [0.01-0.03] |
| *C\** | 4 | 1,895 (256) | 395 (19) | 348 (18) | 47 (1) | $5.5 \times 10^{-8}$ [5.0-6.1$\times 10^{-8}$] | 0.12 [0.09-0.16] |
| *LC\** | 4 | 230 (897) | 502 (627) | 484 (599) | 18<br>(28) | $5.5 \times 10^{-7}$ [5.2-6.3$\times 10^{-7}$] | 0.04 [0.02-0.06] |

<sup>a</sup> The label MMR-<sup>168</sup> corresponds to the aggregation of data from the three MMR-deficient strains constructed from $R^{168}$ ( $\Delta L$ , $\Delta S$ , *L\**). Data for strains $R^{3610}$ and $\Delta S^{3610}$ were obtained from Sung et al. (2015) and mapped to the $R^{168}$ genome sequence.
<sup>b</sup> Conversion between MA-steps and generation based on an estimated average number of generations per step: 25.6 here, 27.53 in Sung et al. (2015).
<sup>c</sup> Number of substitutions in the reference subset of positions well covered by the sequencing data (3,795 kbp). Between parentheses: number of generations or substitutions in time intervals after a detected decrease in mutation rate (discarded from analysis).
<sup>d</sup> Number of transitions. <sup>e</sup> Number of transversions.

Between 113 and 157 nucleotide substitutions were identified after 21 MA-steps in each of the Δ*L*, Δ*S* and *L*\* strains, resulting in point estimates of the substitution rates between 1.4×10^-8^ and 1.9×10^-8^ bp^-1^.gen^-1^ (**Table 1**). These rates were not statistically significant from each other which led us to aggregate the data collected for these three strains under the label MMR-^168^ (**Table 1**). Sequencing an intermediate time-point located at the end of MA-step 11 for the *L*\* strain did not reveal any difference in the rates of accumulation between the first and second parts of the evolution ( **Figure 2** and **Figure S6**). This suggests that in MMR-deficient strains, substitutions identified at endpoints of the MA-lines result from accumulation at a constant rate.

**Figure 2.**
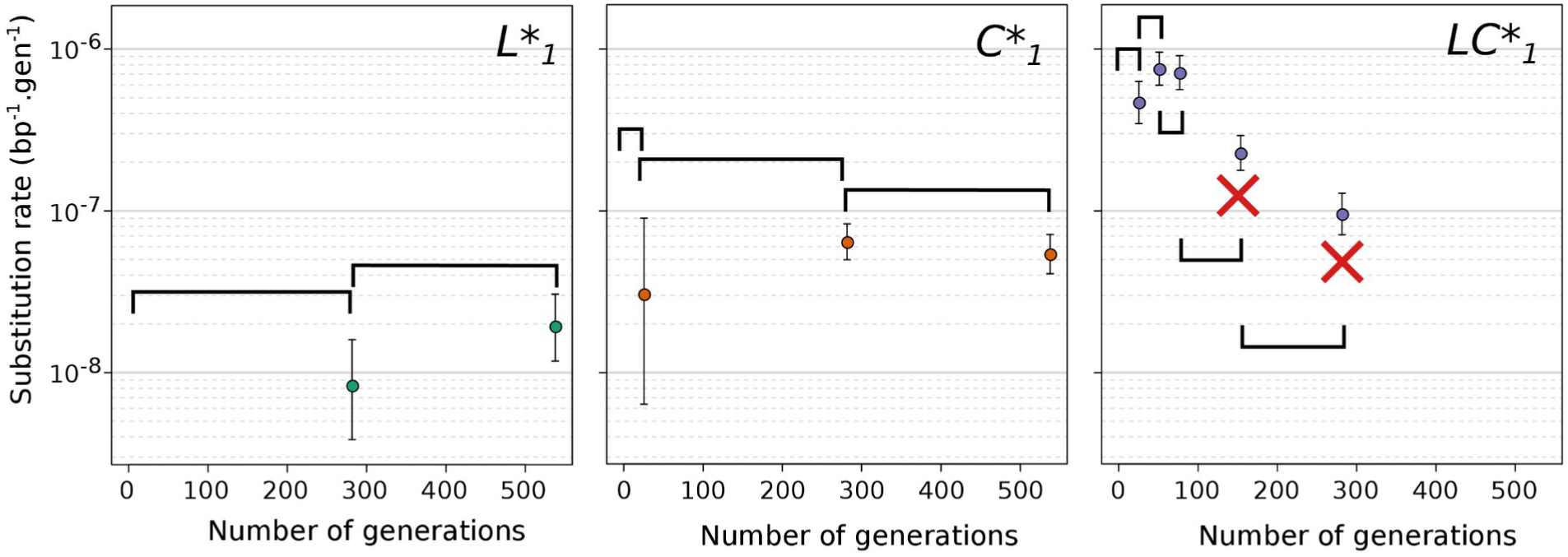
Evolution of substitution rate along mutation accumulation lines. Examples are shown for one MA line from each strain with an inducible mutation rate. The rate is calculated from the number of new substitutions identified within each interval. Sequencing intervals are represented by horizontal brackets and 95% confidence intervals for estimated rates are reported by vertical bars. Red crosses indicate sequencing intervals with significantly decreased mutation rates discarded from downstream analyses.

In contrast to *L*\* MA-lines, both *C*\* and *LC*\* MA-lines showed a tendency towards decreasing substitution rates during their evolution, with differences between the two strains in terms of frequency, temporality, and magnitude of decrease (**Figure 2** and **Figure S6**). A statistically significant decrease was detected for a single *C*\* MA-line (C*_3_) and this decrease was only detected during the second half of the 21 MA-steps evolution. In contrast, a decrease was detected in all *LC*\* MA-lines, as early as during the second MA-step for *LC*\*2 (*p*-value = 8.4×10^-3^), and during MA-steps 3-6 for the three other *LC*\* MA-lines. The magnitude of the decrease was a factor approximately 2.5x for *C*\*3 but reached approximately 50x for *LC*\*4. Therefore, despite heterogeneity between MA-lines, decreases were globally more frequent, quicker and of larger magnitude for *LC*\* than for *C*\*. Importantly, in *LC*\* MA-lines, 56% of the 1,129 identified substitutions occurred in intervals affected by a decrease in the substitution rate (**Table 1**). To determine the mutation rates of the strains, we only considered data from time intervals before any detected decrease. This led to point estimates of the substitution rates for the *C*\* and *LC*\* strains of 5.5×10^-8^ and 5.5×10^-7^ bp^-1^.gen^-1^, respectively (**Table 1**).

Statistically significant decreases in indel rate were also observed (**Figure S7**) and correlated with decreases in substitution rate (**Figure S8**). This is consistent with both proofreading and MMR contributing to correction of indel errors made by PolC polymerase activity (**Supplementary Methods and Results 1.1**), the rate of which increases with homopolymer length (**Figure S9**). The calculated insertions and deletions rates are given in **Table S4**.

Nonsynonymous mutations were found in the inducible synthetic circuits of 4 out of 5 MA-lines exhibiting a decrease in mutation rate (**Table S5**, **Figure S10, Supplementary Methods and Results 1.2**). Given the total number of mutations in the *LC\** lines and the size of the *polC* gene, the number of mutations found on the *polC\** allele is four times higher than expected in the absence of selection (chi-squared test with simulated *p*-values, *p*-value=4.1×10^-2^). This finding echoes previous studies reporting changes in mutation rates in mutation-accumulation experiments (Perfeito et al. 2014; Singh et al. 2017). Indeed, recent studies have concluded that positive selection is possible in mutation-accumulation experiments despite the extreme bottlenecks imposed on the population (Mahilkar et al. 2022; Wahl and Agashe 2022). Here, the over-representation of mutations in the genetic elements that confer the strongest hypermutator phenotypes indicates adaptive evolution through positive selection to reduce the mutation rate. Furthermore, it provides an *a posteriori* experimental justification for the decision to limit mutation-accumulation experiments on proofreading-deficient *E. coli* to 3-6 MA-steps to minimize selection (Niccum et al. 2018).

### Proofreading repairs transversion errors at least as well as transition errors

The sequence data do not provide information about the DNA strand on which the error that led to the mutation originally occurred. To record substitutions, we opted for a framework in which the reference base is the pyrimidine (C or T) of the Watson-Crick pair at the genomic position where a mutation is observed. This allows for a prior-free analysis of strand asymmetries in mutation profiles, and follows the convention used in cancer research (Tate et al. 2019).

In all strains, a slightly higher number of mutations were found on C bases than on T bases (**Figure 3**). All strains also showed a predominance of transitions over transversions, but the strength of this bias differed between strains (**Figure 3** and **Table 1**). The highest proportion of transversions among substitutions, found in R^3610^ (point estimate 0.25), is about 10 times higher than the lowest, found in the MMR-strains (Δ*S*^3610^ and MMR-^168^ strains, point estimates 0.03 and 0.01, respectively). The *C*\* strain showed an intermediate proportion (point estimate 0.12). The small number of transversions made it difficult to compare those changing the reference pyrimidine (C or T) to an A and to a G. Nevertheless, the *C\** and R^3610^ strains may present an excess of C→A over C→G not seen in other strains. The approximately 10-fold increase in the proportion of transitions when comparing the MMR-^168^ strains to the R^3610^ strain is consistent with previously published results on Δ*S*^3610^ (Sung et al. 2015). Indeed, there is general tendency across microorganisms for MMR to reduce the transition rate much more than the transversion rate (Lujan et al. 2012; Long et al. 2018).

**Figure 3.**
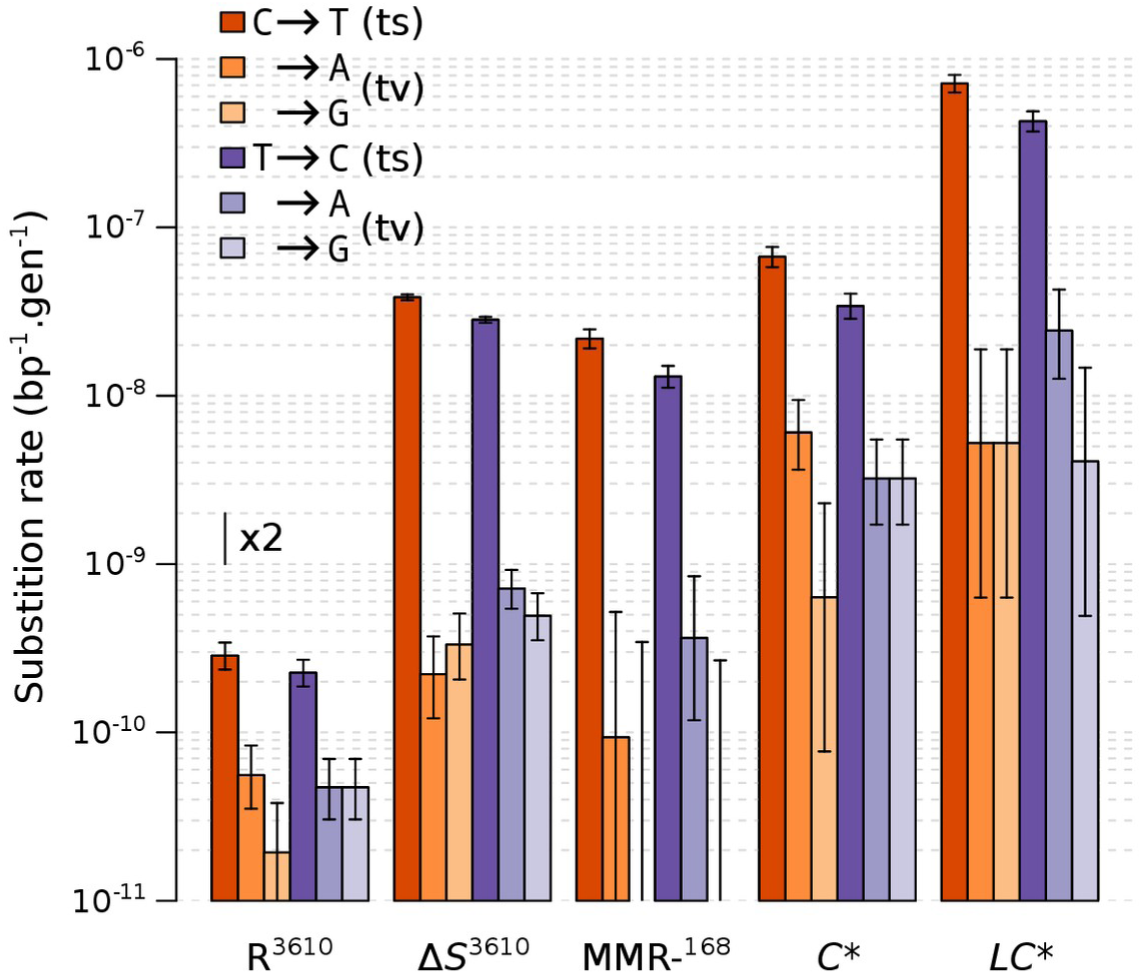
Substitution rates for each type of substitution measured by mutation accumulation experiments. 95% confidence intervals are reported. No C→G and T→G mutations occurred in MMR-^168^, only the upper limit of the 95% CI is shown.

The fold-change in mutation rate was greater when proofreading was inactivated in the presence of MMR than in its absence (**Figure 3** and **Table 1**), otherwise the fold-change in the mutation rate when both repair systems are inactivated compared to the wild-type would be the product of the fold-changes observed when they are inactivated separately (a scenario later referred to as multiplicative fold-changes). This suggests that the efficiency of correction by MMR decreases simultaneously with proofreading inactivation (i.e. in *C\** compared to R^3610^ or R^168^). In *E. coli*, the fold-change in substitution rate upon inactivation of PolIII holoenzyme proofreading was also reported to be much lower in MMR-deficient cells than in wild-type (Schaaper 1988; Niccum et al. 2018). This was interpreted as a consequence of MMR saturation due to the high number of errors introduced during DNA replication, a hypothesis further supported by direct assays of MMR activity and restoration by MMR overexpression (Schaaper 1988; Schaaper and Radman 1989). Comparing *C\** to R^3610^, inactivation of proofreading significantly reduced the proportion of transversions among substitutions (from 0.25 to 0.12), which may also result from MMR saturation in *C\**. Below, we will formally explore the MMR saturation hypothesis using a model-based analysis that integrate information about the chromosomal context of the mutations (adjacent nucleotides and strand).

In the absence of MMR, inactivation of PolC proofreading resulted in a slight increase in the proportion of transversions (from 0.01 in MMR-^168^ to 0.04 in *LC*\*). This difference is not statistically significant (**Table 1**) but suggests that proofreading corrects errors leading to transversions with at least as much efficiency as those leading to transitions.

### Strand-asymmetry of substitution rate at C:G sites is only visible after proofreading

To further characterize the two DNA repair systems, we examined for each strain how substitution rates varied with distance from the origin of replication, strand orientation relative to replication or transcription, coding or non-coding status, and transcription level. For these analyses, we counted substitutions in the different chromosomal contexts and calculated the corresponding “local” substitution rates.

In MMR-^168^ strains, as in Δ*S*^3610^ and R^3610^ strains, substitution rates were significantly higher when C is on the leading than on the lagging strand (**Figure 4A**). This agrees with previous results in *B. subtilis* (Sung et al. 2015), where all mutations were recorded as a change on the “strand templating the leading strand” (*i.e.* the lagging strand), and which showed higher mutation rates for G bases in R^3610^ and Δ*S*^3610^ strains. This bias was not present in the *C*\* and *LC*\* strains, which are proofreading deficient. In addition to the strong asymmetry at C:G sites, we detected a weaker but statistically significant replication-oriented asymmetry in substitutions at T:A sites: the substitution rate is higher when T is on the lagging strand, the difference between the two strands being statistically significant for all our hypermutator strains (**Figure 4A**). In R^3610^, this bias at T:A sites appears to be less pronounced and may even be absent. Consistent with the conclusions of Schroeder et al. (2016), analysis of localization with respect to transcription, which is most often collinear with replication in *B. subtilis*, did not indicate a contribution of transcription-related processes to these asymmetries between strands in any of the hypermutator strains (**Supplementary Methods and Results 1.3, Figure S11AB**).

**Figure 4.**
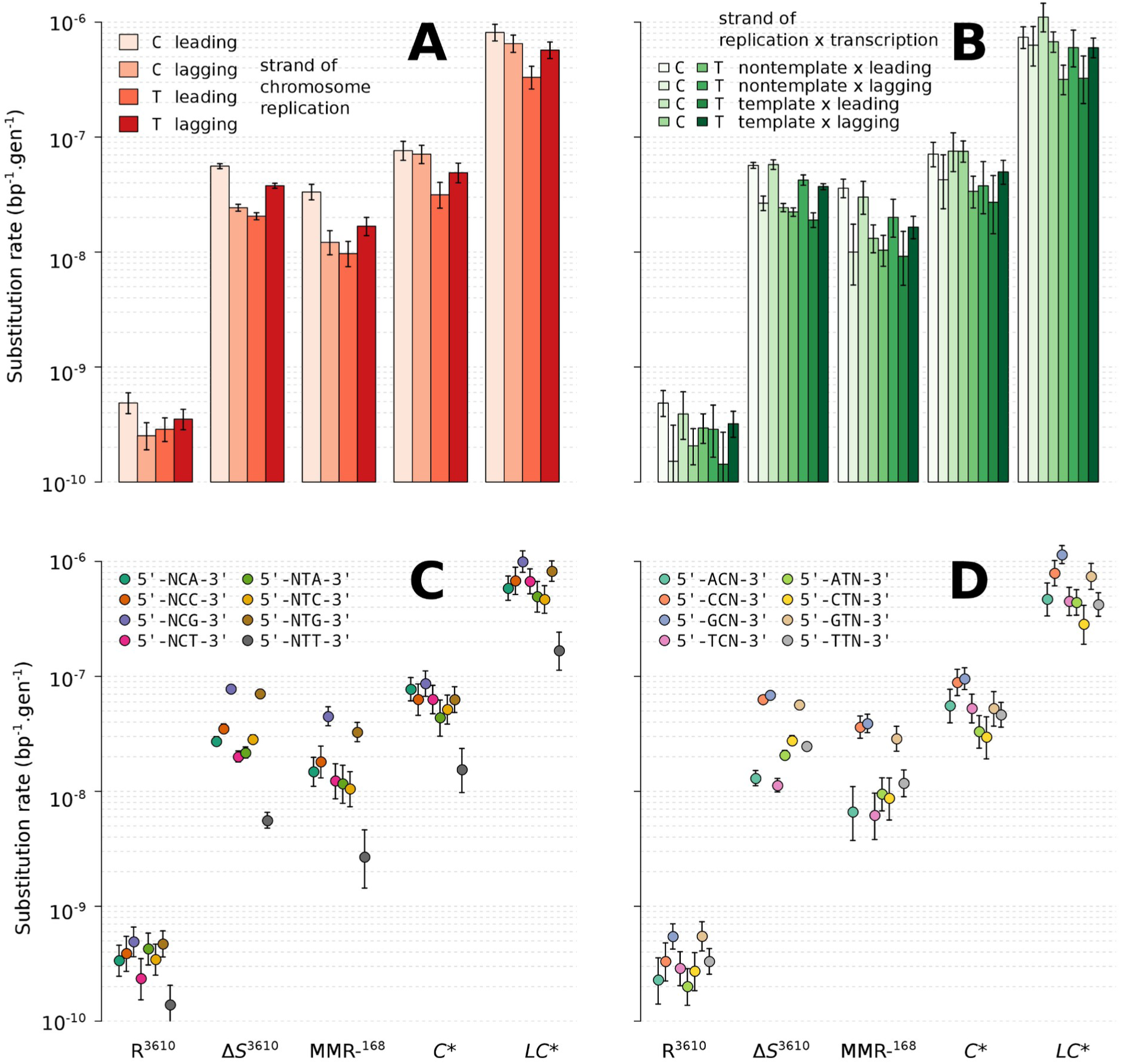
Substitution rate as a function of replication strand and neighboring nucleotides. The pyrimidine of the pair determines the strand of a mutation site. Rates are plotted for each pyrimidine and each genotype. Error bars represent the 95% confidence intervals. **A.** Effect of orientation with respect to the replication strand. **B.** Effect of orientation both with respect to both the replication and transcription strands. **C.** Effect of the 3’-adjacent nucleotide. **D.** Effect of the 5’-adjacent nucleotide.

In wild-type, the replication-oriented asymmetry of substitution rates at G:C sites is a prominent feature of the mutation profile. Consistent with previous analyses of MMR-deficient strains, including Δ*S*^3610^, this bias was also detected in MMR-^168^ strains. Our data revealed its absence in *C\** and *LC\** strains. A similar observation was made in *E. coli* proofreading-deficient strains, leading to the interpretation that proofreading is strand-biased and produces this bias (Niccum et al. 2018), but a concurrent explanation will be discussed below. In contrast, the wild-type had little or no asymmetry at A:T sites, whereas the hypermutator strains have such asymmetry; error correction systems, and in particular the MMR, tend to eliminate this asymmetry.

In parallel, it is intriguing to observe the presence of two trends detected only in wild-type: a higher substitution rate in non-coding regions (**Figure S11C**) and, to a lesser extent, in the half-chromosome near the replication origin (**Figure S11D**). A wild-type specific higher substitution rate in non-coding than coding regions, specific to the wild-type has also been reported in *E. coli* and was interpreted as an indication that “MMR preferentially repairs coding sequences” (Lee et al. 2012; Foster et al. 2018). Alternatively, trends observed exclusively in the wild-type may correspond to substitutions originating from processes not subject to correction by proofreading and MMR, which could be masked by increased substitution rates in hypermutator strains.

### Polymerase initial nucleotide selectivity shapes the distribution of mutations even after proofreading

In all strains, the substitution rate was strongly influenced by the nucleotide adjacent to the focal pyrimidine (**Figure 4C** and **Figure 4D**). This observation, previously made in wild-type and MMR-deficient strains (Sung et al. 2015), extends to proofreading-deficient strains. Simultaneously considering the adjacent nucleotides on both sides (**Table S6** and **Figure S12**) and the replication strand requires binning the counts into 64 replication-stranded triplets. To mitigate the dimensionality problem, exemplified by the absence of observed substitutions for some bins, we adopted a Bayesian estimation framework that incorporates the mean and standard deviation of log-transformed rates as strain-specific hyperparameters (**Supplementary Methods and Results 1.4**). Information from the entire distribution is thereby used to establish point estimates and credibility intervals of the substitution rate for a given triplet (**Figure 5A**).

**Figure 5.**
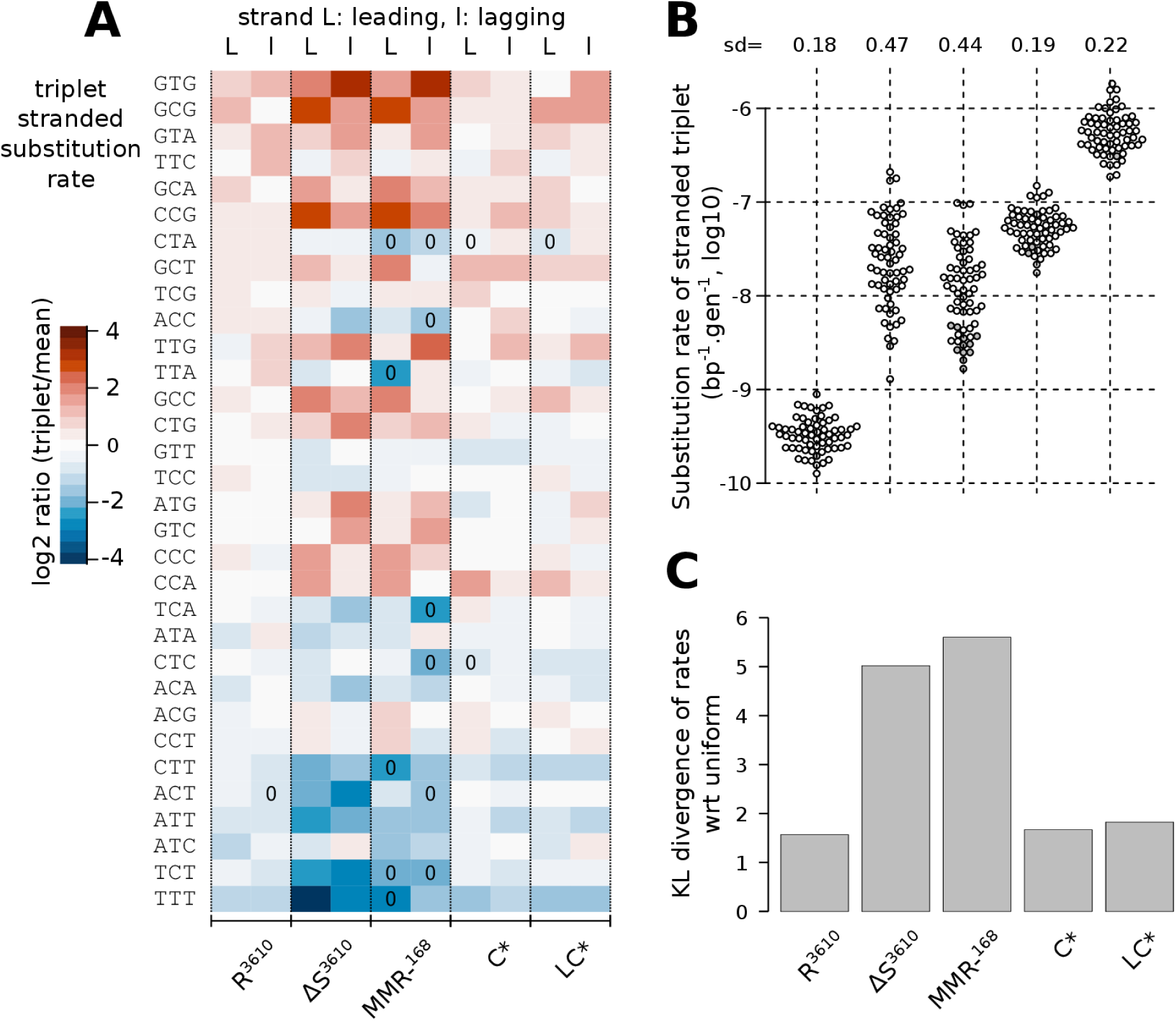
Comparison of stranded-triplet substitution profiles between genotypes. Each stranded triplet corresponds to the pyrimidine of the pair and its 5’ and 3’ nucleotides, distinguishing pyrimidines on the leading and lagging strands of replication. Δ*S*, Δ*L*, and *L\** (IPTG 100 µM) are aggregated as MMR-^168^. **A.** Heatmap representation of the stranded-triplet substitution profiles (log_2_ ratio of the estimated rate with respect to the mean for the genotype) in the different strains. Rates were estimated using a Bayesian method involving a log-normal prior and hyperparameters. Estimates based on the absence of substitutions in this stranded-triplet context are indicated by “0” in the cells of the heatmaps. Triplets are ordered in decreasing order of non-stranded empirical substitution rates. **B.** Beeswarm representations of the distributions of Bayesian estimates of stranded-triplet substitution rates, with the standard deviation of the estimates in log10-scale reported above. **C.** KL divergences from a uniform distribution (same rate for each triplet), derived from robust entropy estimates.

Based on these estimates, we measured a very strong correlation between the stranded-triplet substitution profiles of MMR-^168^ and Δ*S*^3610^ (Pearson correlation r=0.91 for log-transformed rates), confirming that our MMR-^168^ background is very close to the previously studied mutant (Sung et al. 2015). There is also a strong correlation between the stranded-triplet substitution profiles of *LC*\* and proofreading-proficient MMR-deficient strains (r=0.79 between *LC*\* and Δ*S*^3610^). This is consistent with the idea that, after proofreading and before correction by MMR, most of the errors leading to substitutions are due to misincorporation by the PolC polymerase that escaped proofreading. Since MMR removes ∼98-99% of these errors (**Table 1**), they also represent the majority of errors corrected by the MMR.

We also found correlations between the wild-type stranded-triplet substitution profile and those of all hypermutator strains. The correlations are highest with the proofreading-proficient MMR-deficient strains (r=0.72 with Δ*S*^3610^, r=0.71 with MMR-^168^) and remain statistically significant with the proofreading-deficient strains (r=0.58 with *LC*\* and r=0.57 with *C*\*, *p*-values<10^-6^). The correlation between the stranded-triplet substitution rates profiles in *LC*\* and the wild-type, whose global substitution rate is 1600 times lower (**Table 1**), fits remarkably with the working hypothesis that PolC misincorporations that escaped proofreading and MMR substantially shape the substitution profile of the wild-type.

### Proofreading squares the biases of the initial polymerase selectivity

As measured by the standard deviation and the span of their distributions, point estimates of stranded-triplet substitution rates in MMR-^168^ and Δ*S*^3610^ are approximately 10-fold more dispersed than in other strains, doubling the dispersion on a logarithmic scale (**Figure 5B)**. Because point estimates deduced from small counts are not precise, we sought an alternative approach that could directly quantify dispersion. Using a coincidence-counting method for entropy estimation developed to be robust to sparse sampling (Nemenman et al. 2001), we calculated the Kullback-Leibler (KL) divergence between the unknown underlying distribution of observed counts and the theoretical distribution assuming a uniform substitution rate (expected count proportional to the number of occurrences of the triplet in the chromosome). Estimates of KL divergences (**Figure 5C**) confirmed both the similar level of dispersion of substitution rates in the wild-type (R^3610^) and proofreading-deficient strains, and the comparatively much higher dispersion in the MMR-deficient strains (approximately 2.5-fold higher divergence from uniform). In other words, proofreading disperses substitution rates, while MMR activity counteracts this dispersion. This observation is consistent with the conclusions of a study on proofreading of the *E. coli* PolIII holoenzyme (Niccum et al. 2018), and has been interpreted as suggesting a compensation of proofreading biases by opposite biases of MMR correction; a concurrent hypothesis to explain the apparent compensation of the biases is presented in the discussion.

Interestingly, after proofreading (but before MMR correction), the stranded-triplet substitution rates become more dispersed, but remains very similar in terms of the direction of the biases compared to before proofreading (**Figure 5A**). This observation is puzzling because it implies that proofreading amplifies biases that already arise from the sole polymerase activity, rather than simply masking the initial biases with its own biases of greater amplitude. To better understand the implications of this observation, we can formulate a minimal model in which the proofreading activity reduces the initial error probability of the polymerase, denoted γ[i], through a two-step process: first, the detection and removal of a misincorporated nucleotide with probability d[i]; second, the reincorporation of a nucleotide with the error probability γ[i] characteristic of the initial polymerase activity. The error probability after proofreading writes then e[i]=γ[i](1-d[i])/(1-γ[i]d[i]), where the term (1-γ[i]d[i]) accounts for the possibility of cycling the two-step process if a new incorporation error follows the removal. If d[i] is the same for all i, the possibility of cycling, by itself, already amplifies the initial biases. However, this effect is negligible as long as γ[i], which corresponds to the substitution rate observed in the *LC*\* strain (<10^-5^ in **Figure 5B**), remains extremely small compared to 1. Doubling the biases on a logarithmic scale (*i.e.* e[i]/e[j]=(γ[i]/γ[j])^2^), as approximately observed in our data, implies a probability of non-detection and removal in the first step of proofreading (1-d[i]) proportional to γ[i], the error probability of the single polymerase activity. The most common errors made during initial incorporation also the least likely to be corrected during proofreading.

### A theoretical model suggests that MMR can prevent up to 4 mutations per generation before saturation

The apparent efficiencies of MMR and proofreading are strongly influenced by the presence or absence of the other system. As evoked above, proofreading reduced the total substitution rate by a factor 162 in MMR-proficient cells (μ_R3610_/μ_C*_, where μ is the mutation rate under consideration and using the ML estimates from **Table 1**) but only by a factor 32 in MMR-deficient cells (μ_MMR-168_/μ_LC*_). Similarly, MMR reduced the substitution rate more in the presence than in the absence of proofreading. According to Bayesian estimates of substitution rates, none of the triplets or mutation types (ts, tv, ins, del) clearly contradicts this observation (with a posterior probability of being above the diagonal greater than 5% for all triplets; **Figure 6**). The trend becomes even more pronounced when Δ*S*^3610^ is used instead of MMR-^168^ to represent the MMR-deficient proofreading-proficient background (**Figure S12**).

**Figure 6.**
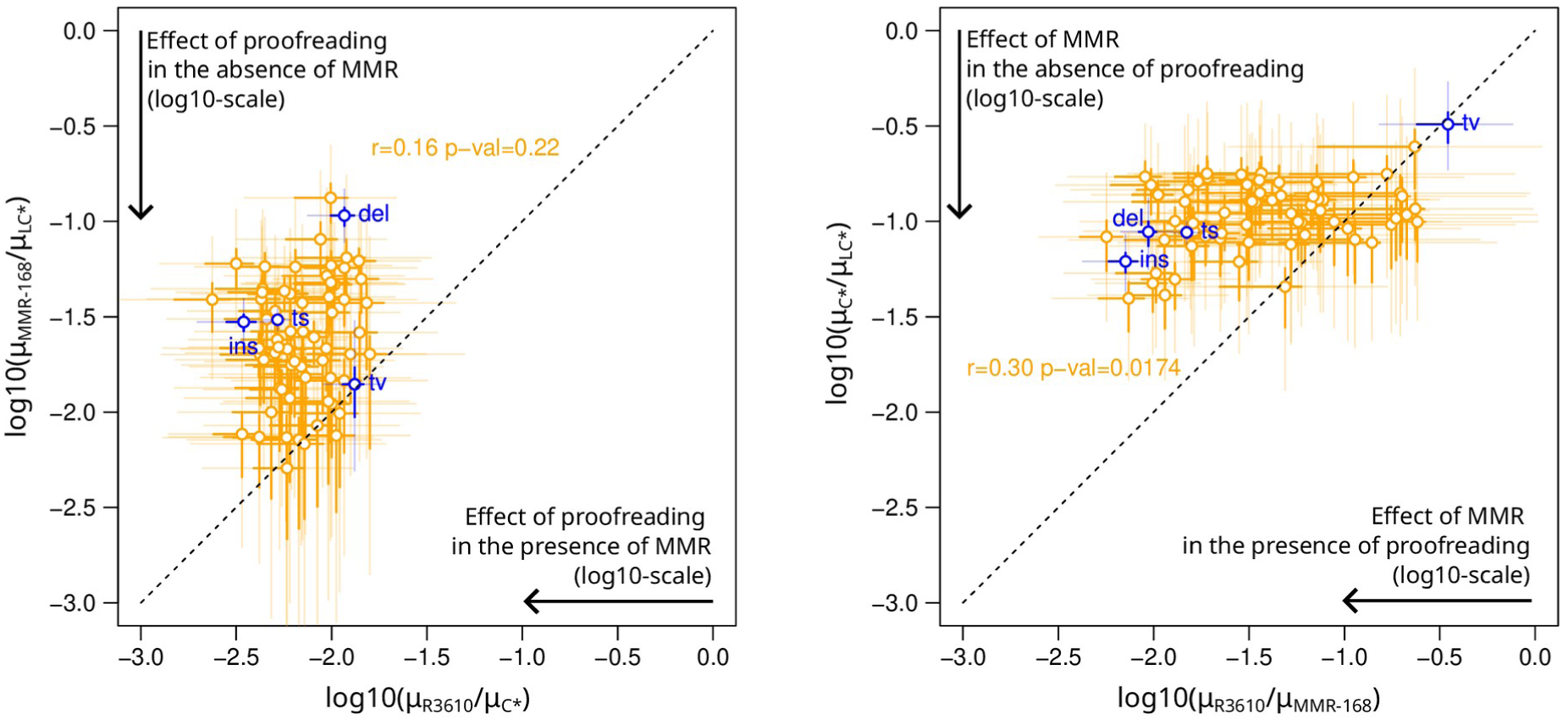
Effects of proofreading and MMR in the presence or absence of the other system. Effects are measured as the log10-ratio of substitution rates. Left plot: effect of proofreading in the presence or absence of MMR. Right plot: effect of MMR in the presence or absence of proofreading. Around each point, the 50% and 95% marginal credibility intervals on horizontal and vertical axes, computed from the quantiles of the posterior distributions, are represented by segments (bold and dark *vs.* thin and light, respectively). The data used for the MMR-substitution profile is MMR-^168^.

To thoroughly investigate the compatibility of the data collected in *B. subtilis* with a saturation mechanism, we formulated a mathematical model in which the mutation rate, μ_C*_[i], in strain *C*\* for a given triplet or type of mutation i is determined by the equation:

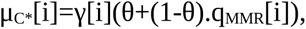

where γ[i] is the error rate before correction by proofreading or MMR, and θ is a mixture parameter common to all values of i. It corresponds to the proportion of errors made by the proofreading-deficient PolC that occur in a physiological context of MMR saturation (generating mutations distributed as in *LC*\*). The complementary proportion (1-θ) is subject to correction by MMR, which reduces the number of errors by a factor qMMR[i]. In this model, whose assumptions are presented along with an associated Bayesian estimation procedure in **Supplementary Methods and Results 1.5**, the mutation rate in *C*\* can be expressed as a function of the rates in the three other backgrounds by identifying q_MMR_[i] with μ_wt_[i]/μ_MMR-_[i] and γ[i] with μ_LC*_[i].

This parameterization was used to estimate the mixture parameter θ and to check the agreement of the model with the experimental data (Figure 7). In practice, the posterior distribution of θ was estimated using either the substitution rates for the 64 replication-oriented triplets (with MMR-^168^ or Δ*S*^3610^ to represent the MMR-deficient background) or the rates of the 4 types of mutations (ts, tv, ins, del). These three posterior distributions are very similar (Figure 7A), with the posterior mean for θ varying only from 0.071 to 0.084. The observed counts, aggregated by mutation type (Figure 7C) and by triplet (**Figures S14** and **S15**), fall within the prediction intervals of the model, indicating a good fit to the experimental data. Notably, although the fraction of errors made by the proofreading-deficient PolC in the context of MMR saturation is small (θ<10%), these errors account for the majority of the mutations observed in strain *C*\* (Figure 7A).

**Figure 7.**
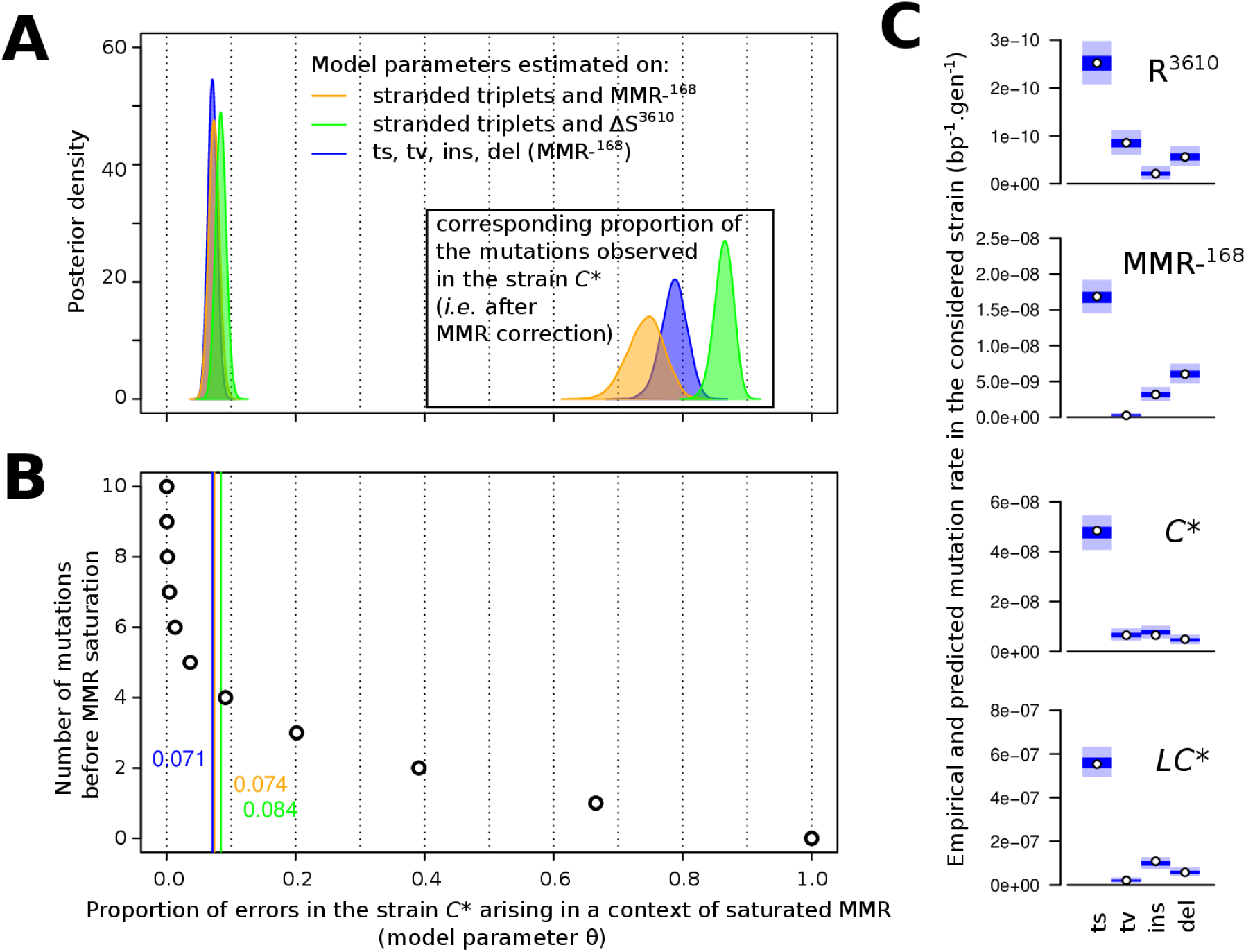
Parameter estimation of the MMR-saturation model and evaluation of the fit to experimental data. **A.** Posterior distribution of the mixing parameter θ corresponding to the proportion of polymerase errors in strain *C*\* arising in the context of saturated MMR. The corresponding proportion of mutations observed in *C*\* (i.e. after MMR correction) is shown in the inset plot. **B.** Relationship between the mixing parameter θ and the number of mutations before MMR saturation in a simplified model of replication: one replication per generation, the first mutations are subjected to MMR correction until MMR saturation. **C.** Evaluation of the fit of the MMR-saturation model to experimentally measured rates of transition, transversion, insertion, and deletion. Points represent empirically calculated mutation rates, *i.e.* the number of observed mutations divided by the number of possible sites in the genome and the number of generations. Colored areas represent the distribution of values for the empirical rates simulated under the posterior distribution of the model parameters (50% of the density in the dark area, 95% including also the light area).

In a simple mechanistic model, these values of θ may correspond to the capacity of the MMR to prevent (with a probability of failure q_MMR_[i]) up to 4 mutations per generation (Figure 7B). Alternatively, θ values can be interpreted as reflecting the fraction of errors made by the proofreading-deficient polymerase before it travels a certain distance after a first error (**Supplementary Methods and Results 1.5**).

## Discussion

### Aggregating mutations in counts: a necessary evil

In principle, our experimental data could also be explained by a model that does not involve MMR saturation. This alternative model recognizes that counting mutations on the genome implies aggregating them by type or context, which are likely to encompass subclasses of mutations arising from errors that are not corrected with the same probability of success. In the extreme scenario where MMR and proofreading correct non-overlapping sets of errors, the increments observed in mutation rates when each system is inactivated would simply add up in cells where both systems are inactivated. This idea has already been mentioned and refuted for mutational signatures in human cancers (Haradhvala et al. 2018). In fact, while changes in mutation rates associated with each repair system are not multiplicative, they are clearly more than additive. This raises interest in a more general model that considers aggregated subclasses of mutations in the counts and arbitrary correction rates. We have algebraically explored this model in its simplest form with only two subclasses (**Supplementary Methods and Results 1.6**). Interestingly, when the probability of error correction differs between subclasses for both MMR and proofreading, the aggregation creates apparent epistasis in the sense that the effect of inactivating one system, as measured by the mutation rate fold-change, depends on the presence or absence of the other system.

With more parameters than data points (5 parameters for 4 mutation rates), even the simplest two-subclass scenario can fit almost any data set, explaining additive to super-multiplicative effects. The model is therefore difficult to falsify. We note, however, that the sign of this epistasis depends on whether the systems have similar or opposite specificities: if they tend to repair the same subclasses of errors the apparent epistasis will be positive (super-multiplicative), otherwise the epistasis will be negative (sub-multiplicative, as in our data). Explaining negative epistasis with this model is therefore at odds with the widely accepted idea that MMR correction is mostly coreplicative and targets the same errors as those corrected by proofreading, *i.e.* Watson-Crick mismatches introduced by DNA polymerase during DNA replication. If subclass aggregation does not contribute significantly to the observed apparent epistasis, which is well explained by MMR saturation, it still complicates the interpretation of the apparent efficiency of repair systems deduced from mutation profiles.

### A cautious interpretation of the apparent efficiency of proofreading and MMR

The proofreading-deficient strain *C*\* should not be considered as fully MMR-proficient due to MMR saturation. Therefore, its mutation rate cannot be used to measure the apparent efficiencies of proofreading and MMR under wild-type physiological conditions, i.e. as the numerator in a ratio μC*/μLC* to estimate the probability that an error escapes proofreading, or as the denominator in a ratio μwt/μC* to estimate the probability that an error escapes MMR correction. Instead, these repair system escape probabilities should be estimated from the ratios μ_MMR-_-/μ_LC*_ and μ_wt_/μ_MMR-_- shown in Figure 8 (where MMR-is either MMR-^168^ or Δ*S*^3610^) for each replication-oriented triplet and mutation type (ts, tv, ins, del). The rates of errors leading to mutation before and after correction by the combined action of proofreading and MMR (mutation rates in *LC\** and wild-type, respectively) are also shown.

**Figure 8.**
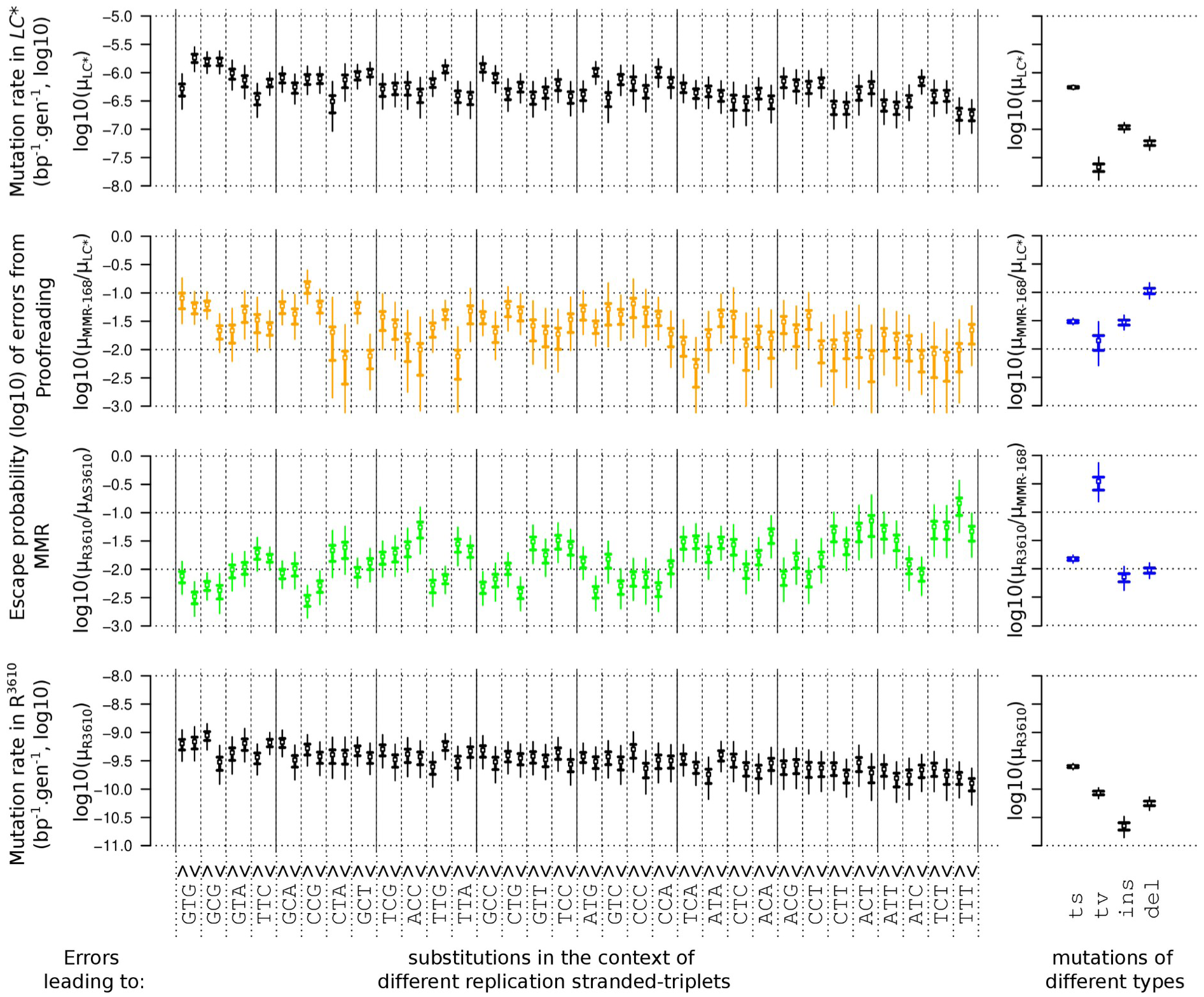
Apparent efficiency of proofreading and MMR across replication-stranded triplets and mutation types. Estimated proofreading and MMR escape probabilities are shown (middle plots) along with estimated mutation rates in the absence or presence of both correction systems (top and bottom plots). Replication-stranded triplets are ordered in decreasing order of non-stranded empirical substitution rates and then focal pyrimidine on the leading and lagging strands of replication. Bold and thin vertical bars represent 50% and 95% credibility intervals, respectively.

Proofreading and MMR escape probabilities (μ_MMR-_-/μ_LC*_ and μ_wt_/μ_MMR-_) are correlated with triplet substitution rates (**Figure S16**). In Figure 8, the triplets are ordered according to wild-type substitution rates (as in Figure 5A), which highlights general trends: proofreading escape tends to be higher on triplets where the initial polymerase selectivity is the lowest (**Figure S16**, r=0.32 *p*-value=0.0097); MMR escape tends to be lower on triplets with high wild-type mutation rate (**Figure S16**, r=-0.45, *p*-value=0.00018). Note that these two correlations are of opposite sign to the artifactual correlations that would be generated by noise in the estimates of μ_LC*_[i] and μ_wt_[i], which also enter in the ratios μ_MMR-_[i]/μLC*[i] and μ_wt_[i]/μ_MMR-_[i].

The positive correlation between proofreading escape and mutation rate in *LC\** is consistent with the observation that proofreading increases the bias of polymerase errors. Statistical uncertainty about the probability of proofreading escape makes it difficult to draw conclusions for specific replication-stranded triplets, and few triplets show clear strand asymmetry. It is interesting to note, however, that this positive link is not seen within pairs of triplets that differ only by the strand. In particular, G<u>T</u>G and A<u>T</u>G are more affected by substitutions in *LC\** when the focal pyrimidine (T) is on the lagging strand, but no difference in proofreading escape probability between strands is detected. For these triplets, the initial strand asymmetry may originate from incorporation errors made by the DnaE polymerase during synthesis of the lagging strand that are efficiently proofread *in-trans* by PolC. Conversely, errors affecting GCT and GCG appear to be less corrected by proofreading when the C-site is on the leading strand, but the substitution rates in *LC\** are similar between strands. In these cases of C-site substitutions on the leading strand, the apparent probability of proofreading escape most likely overestimates the escape of initial polymerase selectivity errors because of the likely contribution of deamination damage to the mutation profile of the proofreading-proficient MMR-deficient strains.

Deamination of cytosine to uracil in the lagging strand template (*i.e.* the leading strand) due to exposure of single-stranded DNA in the context of the replication fork has long been suspected to be the reason for the near-universal asymmetry between replication strands in prokaryotes (Frank and Lobry 1999) and indeed has been shown to be a substantial source of C→T transitions in wild-type (Bhagwat et al. 2016). Strikingly, proofreading-proficient MMR-deficient strains exhibit a similarly oriented strong strand asymmetry for substitutions at C sites (Lee et al. 2012; Sung et al. 2015). It has been hypothesized that cytosine deamination may also cause this asymmetry (Foster et al. 2018) which is consistent with other studies reporting a role for the *B. subtilis* MMR in counteracting the effects of base deamination (López-Olmos et al. 2012; Patlan-Vazquez et al. 2022). In the absence of proofreading, the high rate of polymerase misincorporation on both strands would mask the deamination-induced asymmetry.

The hypothesis that proofreading efficiency is positively linked to initial polymerase selectivity was proposed above based on the analysis of substitution rates by triplet context without distinguishing between transversions and transitions. Transversions account for only 4% of the substitutions in *LC\**. The lower apparent proofreading escape probability of errors leading to transversions than to transitions (Figure 8) is thus consistent with the above hypothesis. However, the difference between the proofreading escape probabilities of the two types of errors, which cannot be precisely estimated, seems to be smaller than expected. A possible explanation would that the apparent proofreading escape probability overestimates proofreading escape for initial polymerase selectivity errors that lead to transversions. This would occur if a substantial fraction of the rare transversions in proofreading-proficient in MMR-deficient strains did not originate from polymerase errors and thus was simply not subjected to proofreading.

Regarding MMR escape, which is not the primary focus of this study, its strong negative correlation with proofreading escape (**Figure S17**, r=-0.63, *p*-value=2.6×10^-8^) could reflect an evolution of MMR towards correcting the most common DNA replication errors. Indeed, this would echo the proposed role of differential MMR efficiency in balancing DNA replication fidelity between the two strands in eukaryotes (Lujan et al. 2012; Andrianova et al. 2017; Zhou et al. 2021).

However, the negative correlation between the apparent probabilities of MMR and proofreading escape could also reflect an overestimation of MMR escape for the sites least affected by DNA replication errors. Indeed, the contribution of different sources of spontaneous mutations in wild-type remains difficult to characterize (Schroeder et al. 2018), and thus there is a lack of knowledge about the fraction of the wild-type mutation profile that is not subject to MMR correction. Among the studies addressing this question, it has been shown that inactivation of oxidative damage repair pathways increases the mutation rate in *E. coli* under optimal growth conditions without external stress, suggesting that damage that escapes correction by oxidative damage repair pathways may also contribute to the wild-type mutation profile (Foster et al. 2015). Since oxidative damage tends to cause transversions, this hypothesis would be consistent with the higher proportion of transversions in the wild-type than in any of our hypermutator strains. If a substantial fraction of the wild-type substitution profile μ_wt_ originates from sources not subjected to MMR correction, variations not directly related to MMR efficiency may enter the ratio μ_wt_/μ_MMR-_-used to estimate the MMR escape probability (Figure 8), both via its numerator μ_wt_ (*e.g*., variations in the amount of substitutions originating from these other sources) and via its denominator μ_MMR-_- (*e.g*., variations in the amount of polymerase errors remaining after proofreading). Indeed, the very strong negative correlation across triplets between μ_wt_/μ_MMR-_ and μ_MMR-_- (**Figure S16**, r=-0.94, *p*-value=2.3e^-31^) suggests that variations in μ_MMR-_ may be largely responsible for variations in the apparent MMR escape probability-. Combined with the unsurprising and clear positive correlation between the apparent proofreading escape probability (μ_MMR-_-/μ_LC*_) and μ_MMR-_-(**Figure S16**, r=0.71, *p*- value=7.3e^-11^), the substitutions originating from sources not subjected to MMR correction may thus contribute to the observed negative correlation between the apparent probabilities of MMR and proofreading escape.

## Conclusion

Saturation of the MMR in the absence of proofreading, greater dispersion of substitution rates in the presence than in the absence of proofreading, and the existence of strand biases that become apparent only in the presence of proofreading appear to be traits shared between *B. subtilis* and *E. coli*. Given the considerable divergence between these two organisms in terms of phylogenetic distance and molecular organization of DNA polymerase and MMR, these traits are likely to be common to many other organisms. The characterization of the overdispersion of mutation rates in MMR-deficient proofreading-proficient strains compared to other strains led us to conclude that proofreading skews DNA polymerase error rates. This could be interpreted as an inherent drawback of the proofreading principle, which relies on the DNA polymerase to detect its own errors. The role of judge and party leads to the same biases in initial nucleotide selectivity and error correction efficiency.

This study also examines the consequences of aggregating mutations in counts and, more generally, recognizes the difficulty of interpreting the apparent efficiencies of repair systems. First, unaware aggregation of subclasses of mutations resulting from different molecular pathways can create almost any pattern of apparent interactions in the effects of disabling different repair systems. Second, interpretation of the effect of disabling one repair system is generally complex because of uncertainty about the contribution of damage or errors subject to correction by that system to the mutations observed in its presence. It is thus difficult to interpret the appearance of strand asymmetry upon proofreading activation, which may be caused by a post-proofreading mutation process, such as deamination, rather than a proofreading bias. Similarly, the “flattening” of mutation rates upon activation of the MMR is difficult to interpret, as it could result from a better efficiency of the MMR in correcting the most common errors, but also from the contribution of multiple sources to the wild-type mutation profiles. Indeed, a substantial contribution of multiple sources seems consistent with the drift-barrier hypothesis (Lynch et al. 2016): in the wild-type, the rates of mutations resulting from different molecular pathways may have been pushed below the same point where they do not encounter significant counter-selection.

The construction of strains with inducible hypermutator phenotypes addressed the stability issue that affected the reproducibility of some studies on *E. coli* proofreading-deficient strains discussed in (Niccum et al. 2018). It was also motivated by the interest in tools for synthetic biology applications. Regarding the future use of our inducible systems to accelerate evolution in the laboratory, a proofreading-deficient strain would yield a less skewed mutation spectrum than an MMR-deficient strain. If necessary, extreme mutation rates can be achieved by disabling both repair systems simultaneously, but strong counter-selection against mutational load makes such induction feasible only for a short period of time.

## Methods

### Media and bacterial strains

*E. coli* DH5α was used for plasmid construction and transformation using standard techniques (Sambrook et al. 1989). The *B. subtilis* strains used in this study were derived from our Master Strain (MS), a prophage-free and *trp*^+^ derivative of *B. subtilis* 168 (Dervyn et al. 2023), denoted here R^168^. Lysogeny broth (LB) was used to grow *E. coli* and *B. subtilis*. Transformation of *B. subtilis* cells was performed using the protocol of (Konkol et al. 2013). When relevant, the media were supplemented with the following antibiotics: ampicillin 100 μg.mL^-1^ for *E. coli* and spectinomycin 100 µg.mL^-1^ or kanamycin 5 µg.mL^-1^ for *B. subtilis*.

### Construction of hypermutator strains

The Δ*S* and Δ*L* mutant strains were generated by transforming the PCR-amplified *kan mutL* and *mutS kan* sequences using the P1-P2 primer pair together with genomic DNA from Δ*mutS*::*kan* and Δ*mutL*::*kan* mutant strains, respectively. These strains were obtained from the previously published *B. subtilis* single-gene deletion library (Koo et al. 2017).

To construct the *L*\* strain, the first and second halves of the *mutL* gene were PCR-amplified using primer pairs P5-P8 and P6-P7, respectively (**Figure S1**), where P7 and P8 both carry the desired point mutation (as indicated in **Table S7**). The two fragments were then assembled by PCR, resulting in the *mutL*(N34H) allele. The backbone of the pDR111 plasmid (kind gift of D. Rüdner) containing the isopropyl-β-D-1-thiogalactopyranoside (IPTG) inducible P*_hyperspank_* promoter (denoted here P*_hs_*) and the *spec* gene (conferring resistance to spectinomycin), was PCR-amplified using P3 and P4. The 5’ extensions of P5 and P6 then allowed the assembly of the *mutL*(N34H) allele with the PCR-amplified pDR111 using the HiFi DNA assembly protocol (New England Biolabs, USA). This resulted in cloning the *mutL*(N34H) allele under the control of P*_hs_* into a *B. subtilis amyE-* integrative plasmid (**Figure S1**).

Similarly, to construct the *C*\* strain, the *polC* allele found in *B. subtilis mut*-1 (Bazill and Gross 1973), characterized by the G430E and S621N mutations (Sanjanwala and Ganesan 1991), was PCR-amplified using the primer pair P11-P12 and assembled to the PCR-amplified pDR111 (using P3 and P4) using the HiFi DNA assembly protocol (New England Biolabs, USA). This resulted in cloning the *polC mut*-1 allele under the control of P*_hs_*into a *B. subtilis amyE-*integrative plasmid (**Figure S2**).

For the construction of the *LC*\* strain, a *mutL*\* *polC*\* synthetic operon was generated by assembling the *mutL*(N34H) allele PCR-amplified from strain *L*\* using P5 and P11, and the *polC mut*-1 allele PCR-amplified from *C*\* using P12 and P10, to the PCR-amplified pDR111 (using P3 and P4) using the HiFi DNA assembly protocol (New England Biolabs, USA). This resulted in the cloning of the *mutL*\* *polC*\* synthetic operon under the control of P*_hs_* into a *B. subtilis amyE-* integrative plasmid (**Figure S2**).

Plasmids were transformed into the *B. subtilis amyE* locus by double recombination events. All strains were verified by sequencing, and transcriptomics experiments were performed to compare global gene expression. The RNA-seq reads and detailed protocols and results have been deposited in GEO.

### Fluctuation assays

For each strain to be tested, a single colony was grown in 1 mL LB at 37°C for 90 minutes. This preculture was serially diluted in fresh LB to start cultures with a small number of cells (N_0_). Cells were then grown for 7.5 h to reach saturation. When induction with IPTG was tested, LB medium with the desired concentration of IPTG was prepared from an IPTG stock concentration of 1 mM just before use. When the culture volume was 1 mL, the cultures were centrifuged before plating to retain the cells, and 750 µL of supernatant was removed. The remaining 250 µL were gently vortexed before plating onto LB supplemented with rifampicin (10 µg.mL^-1^). For each assay, a number of cultures (8 for the R^168^ assay, 4 for the Δ*S* and Δ*L* assays, and 3 for all other assays) were not plated on LB medium supplemented with rifampicin, but were serially diluted and plated on LB agar to determine the final number of cells (N_t_). Fluctuation assays performed on the same day were considered to have the same distribution of final cell numbers. All other cultures were plated on LB agar with rifampicin to determine the number of Rif-resistant colony-forming units (CFUs). All plates were incubated at 37°C and scored for CFUs after 24 h of growth.

The maximum likelihood estimator (MLE) of the number of mutations per assay (m) and the confidence interval, were calculated under the Luria-Delbrück model taking into account the variation in the final number of cells (Zheng 2016), using the newton.B0 with default parameters and confint.B0 functions of the R package “Rsalvador” v1.7 (Zheng 2017). For the calculation of confidence intervals, the initial guess for the parameter m was taken as the m given by the “newton.B0” function. The mutation probability was assumed to be constant over the cell cycle, so that the mutation rate per base per generation is the mutation rate per base per cell division (Foster 2006). The final number of cells, N_t_, is the result of N_t_ - N_0_ cell divisions, *i.e.* ∼N_t_ divisions. The rate of Rif^R^ emergence was therefore calculated as μ_Rif_= m/N_t_.

### Mutation-accumulation experiments and sequencing

One isolated colony was collected each day (24 h at 37°C), suspended in culture medium + 20% glycerol, and diluted by 2×10^5^, a factor that allows distinguishable colonies, before plating on LB agar (+ 100 µM IPTG for *L*\*, *C\** and *LC\** strains) for the next MA-step. Counting of the colonies present on the agar plate provided an estimate of the number of bacteria initially present in the diluted colony and thus the number of generations per MA-step. Four parallel MA-lines were propagated per strain (21 consecutive MA-steps for Δ*S*, Δ*L*, *L*\* and *C\**, 11 for *LC\**).

For sequencing at intermediate and end points of the MA-lines, 5 to 50% of the picked colony was cultured in LB medium to collect cells. DNA was extracted using the GenEluteTM Bacterial Genomic DNA Kit (Sigma-Aldrich) according to the supplied protocol. DNA samples corresponding to an intermediate time-point in the four parallel MA-lines for the same strain were pooled in equimolar proportions. Individual and pooled DNA samples were sequenced (150 bp paired-end reads) on an Illumina platform (NovaSeq 6000) to an average depth of ∼300. Reads are available in NCBI SRA.

### Detection of mutations

The reads were aligned to the reference sequence of the *B. subtilis* 168 genome (GenBank: AL009126.3) using BWA-MEM v0.7.17 (Li and Durbin 2009), after quality control and trimming using Sickle v1.33 (command “sickle pe” with options “-t sanger -x -q 20 -l 20”) (Joshi and Fass 2011). Properly paired reads, selected using “samtools view -f 3” (samtools v1.14, Li 2011), were locally realigned around indels using ABRA2 v2.24 (Mose et al. 2019). The number of occurrences of each nucleotide (base read quality ≥35) and indel at each position of the reference in confidently mapped reads (alignment quality ≥50) was counted using “samtools mpileup” with the options “-aa -d 5000 -q 50 -Q 35 -x -B”. These counts were analyzed in R.

For each position, the effective sequencing depth (DPeff) was calculated as the total number of informative reads. For the computation of the mutation rates, a reference subset of positions common to all samples was determined. This reference consisted of positions that were well covered on both strands (DPeff≥100 and ≥10% of the reads on the less represented strand) in all samples. Most of the regions with low coverage corresponded to the regions deleted in the construction of the MS / R^168^ strain (Dervyn et al. 2023), which lacks 233.4 kb of chromosome relative to AL009126.3, and to the multicopy structural RNAs. Over-covered regions were also eliminated from this reference subset of positions and consisted of: the region of gene *upp* and downstream (positions 3,788,426 to 3,789,124), which was repeated due to pop-ins and pop-outs at this locus during the construction of the R^168^ strain; the region from position 2,432,478 to 2,433,315, over-covered in *polC\** samples; the regions of the genes *polC* (1,727,133 to 1,731,446) and *mutL* (1,778,337 to 1,780,539) duplicated by insertion of the mutant alleles *polC\** and *mutL\**. This resulted in a reference subset of 3,794,734 positions (out of a total of 4,215,606 bp in AL009126.3), which served as our reference chromosome for the mutation rate calculations.

The distribution of the proportion of non-reference reads in the different samples was examined graphically to establish relevant cut-offs for the identification of mutations. A mutation was identified at the endpoint of an MA-line if a variant accounted for ≥75% of the DPeff at a position, with ≥10% of the non-reference reads on the less represented strand. If intermediate time-points were available for this MA-line, the mutation was traced back to the first time-point where it occurred at frequency ≥5% in the corresponding pooled sequence sample. Due to the detection, during graphical examination, of contamination from other samples, we lowered the cut-off from 75% to 60% for the identification of mutations in the third MA-line of Δ*S* and from 5% to 2% for the analysis of the pool corresponding to the intermediate time-point for *L*\* strain MA-lines. Mutations found in all samples or in the four MA-lines of a same strain were interpreted as fixed before the mutation-accumulation experiment and were discarded for the calculation of mutation rates.

### Detection of mutations in inducible synthetic circuits

The reads were also aligned to the reference sequences of the inserted regions represented in **Figure S2**. For the mutant alleles *mutL*\* and *polC\**, reads from the native alleles (*mutL* and *polC*) were also mapped to the insert, and variant calling *per se* cannot distinguish between a mutation in the native and mutant allele. However, at positions where bases differed from the reference on these genes in individual samples, the proportion of these alternative bases was bimodally distributed, with two peaks, located at approximately at 40% and 60% of the reads **(Figure S10**). Since the characteristic point mutations of both *polC\** and *mutL*\* accounted for more than 50% of the reads ( between 52% and 71% and between 65% and 75%,, respectively) at their respective positions, we predicted that the mutations for which the majority of the reads differed from the reference would be on the mutant allele, while the others would be on the native allele. This prediction is consistent with the chromosomal position of the *amyE* locus where the mutant alleles are inserted, *i.e.* closer to the origin of replication of the chromosome than either native allele, and thus expected to be more abundant in the sample due to ongoing replication. To verify this, we used specific primers to amplify and sequence either the native or the inserted allele. Of the 5 PCR-verified mutations, all were found on the predicted allele (**Table S5**).

### Chromosome partitioning to assess the effect of transcription and replication on mutation rate

To assess the effect of transcription, the “gene” features of the GenBank annotation served to define the dichotomy between “template” and “nontemplate” strand as well as between “coding” and “noncoding”. Since the “noncoding” represents only ∼10% of the genome and includes transcribed untranslated regions (UTRs) we also sought to assess the impact of transcription with more statistical power and precision than allowed by the GenBank annotation. For this purpose, two categories of regions of approximately equal size were defined based on the transcribed regions identified in 269 samples of a wild-type strain representative of a wide variety of growth conditions (Nicolas et al. 2012). These two categories reflected the amount of transcripts in LB as measured in 9 samples corresponding to growth in liquid LB (triplicate samples for exponential, transition and stationary phases) and 2 samples corresponding to 16 hours of growth on LB agar (non-onfluent colonies). The “high” transcription level regions were those that were in the top 30% in at least one of these 11 samples while the “low” transcription level regions were those that were never in the top 30%. All overlapping regions (*i.e.* both strands were transcribed) were eliminated, as well as all regions shorter than 100 bp. This resulted in a set of 3,622 non-overlapping transcription-oriented regions covering 84.9% of the reference genome (43.4% for “high”, 41.4% for “low”).

To assess the effect of DNA replication asymmetry, the leading and lagging strands were defined based on the origin of replication (position 1) and its most central terminus (position 2,018,289) (Wake 1997). To assess the effect of DNA replication timing, the genome was divided into a “first half” corresponding to the 2 Mbp of the chromosome centered on the origin of replication (position 1) and a “second half”.

### Mutation rate estimations and comparisons

To include the list of mutations in R^3610^ and Δ*S*^3610^ from (Sung et al. 2015) and (Sung et al. 2016) in our analysis, the positions on the *B. subtilis* NCIB 3610 genome (GenBank: CM000488.1) were transferred to the *B. subtilis* 168 genome by mapping the 41 bp long sequence centered on each mutation site. Retaining only exact and unique matches, more than 99% of these mutations were transferred (**Table S3**), with perfect collinearity between the positions of the mutations on both reference genomes.

Maximum-likelihood estimates of mutation rates were obtained as μ=m/(T×G), where *m* is the total number of mutations of a given type in a given genotype and genomic context (nucleotide at focal position and adjacent nucleotides, orientation with respect to replication, transcription, …), T is the total number of occurrences of the genomic context in the reference sequence, and G is the number of generations considered in MA-lines. Confidence intervals for these point estimates were calculated using the exact method for Poisson counts implemented in R package “epitools” v0.5-10.1, with *m* as count and T×G as time-person at risk.

To assess whether a factor affects the substitution rates, we used Generalised Linear Models (GLMs) for Poisson distributed count data with log-link, combined with an Analysis of Variance (ANOVA) (R package “stats” v3.6.3) to compare the fit of a GLM including and a GLM excluding the factor of interest. This statistical comparison was performed separately for each genotype.

Markov chain Monte Carlo methods implemented in JAGS (Plummer 2003), accessed via the R package “rjags”, were used for Bayesian estimation via posterior sampling, in particular for the estimation of mutation rates of replication-stranded triplets and the MMR saturation parameter θ. Models and algorithm settings are described in detail in **Supplementary Methods and Results 1.4**.

### Mathematical modelling of mutation rates

Assumptions and Bayesian estimation procedure for the model with saturation of the MMR are presented in **Supplementary Methods and Results 1.5**. Algebraic analysis of the general model with two subclasses of errors and two repair pathways is presented in **Supplementary Methods and Results 1.6**.

## Supporting information

Supplementary Methods and Results

## Data access

The RNA and DNA sequencing data generated in this study have been submitted to the NCBI Gene Expression Omnibus database under accession number GSE239804 and to the NCBI Sequence Read Archive under BioProject accession number PRJNA995423, respectively.

## Acknowledgements

We express our gratitude to Guillaume Achaz, Elena Bidnenko, Clément Nizak and Paulo Tavares for their advice during the course of this work. We also thank the INRAE MIGALE bioinformatics facility (https://doi.org/10.15454/1.5572390655343293E12) for providing computational resources and Marina Elez for helpful comments on the manuscript.

## Author contributions

IT, MJ, and PN conceived the project and designed the experimental plan. ED and IT performed the experiments. IT processed the raw data with the help of CG. GKKK, IT, and PN conducted the statistical and mathematical analyses. IT, MJ, and PN interpreted the results and wrote the manuscript with contributions from all authors.

