## Supplementary Methods and Results for "The mutational landscape of *Bacillus subtilis* conditional hypermutators shows how proofreading skews DNA polymerase error rates"

Provided as distinct sheets in a separate xlsx file:

- Table S3. List of all substitutions and indels in MA-lines.
- Table S6. Proportion of transversions by replication-oriented triplet.

### 1 Supplementary Methods and Results

#### 1.1 Proofreading and MMR correct the indels introduced by the polymerase activity of PolC

We examined the accumulation of short insertions and deletions (indels of length 1 or 2 bp) in the MA-lines (**Table S5**). Although the small number of indels compared to substitutions makes it more difficult to detect statistically significant changes, the MA-lines that showed the largest decreases in substitution rates ($LC^*_1$ and $LC^*_4$) also showed statistically significant decreases in indel rates (**Figure S8**). As expected, insertion and deletion rates were lower than substitution rates, with ratios of up to 5 (**Table S5** and **Figure S9**). In general, indel rates and substitution rates exhibited a strong positive correlation (Pearson r=0.93, *p*-value=$2.28\times10^{-16}$) across time intervals, MA-lines, and strains (**Figure S9**). This correlation is consistent with a contribution of both MMR and proofreading to the repair of substitutions and indels.

After discarding the same time intervals as for the substitution rates, all short indels except a single deletion were found in homopolymers or tandem repeats of length ≥2. The vast majority of indels involved a single nucleotide (length 1), with only five indels of length 2, all consisting of a dinucleotide motif in a tandem repeat. Overall, 94% of deletions and 97% of insertions occurred in homopolymers of length ≥4. These results are consistent with the model of (Streisinger et al. 1966), according to which displacement of the template strand or newly synthesized strand is more frequent in microsatellite regions, and thus indels are more frequent in these regions; with indel rates increasing with the length of the homopolymers (**Figure S10**). Using a GLM (Poisson data with log-link), we modeled indel rates in the homopolymers with a univariate quadratic function of homopolymer length, where the first-order term captures the constant initial slope between homopolymer lengths 2 and 6 and the second-order term captures the decrease in slope after length 7. The first-order terms were not significantly different between strains *C**, *LC** and $R^{3610}$, indicating that PolC proofreading and MMR activities do not significantly contribute to the increase in indel rate with the homopolymer length, which thus results solely from the PolC polymerase activity. Similar trends have been observed in yeast for these homopolymer lengths of when MMR and/or proofreading are abolished (Lujan et al. 2015).

In the presence of IPTG, the strain *LC**, in which both major repair systems are inactive, is more prone to insertions (66% of insertions among all indels, **Table S5**). In contrast, MMR-$^{168}$ and R$^{3610}$ have significantly lower insertion than deletion rates (only 27% of insertions among indels in R$^{3610}$). This suggests that PolC-mediated DNA replication tends to cause more deletions than insertions, but this bias is reversed in wild-type by PolC proofreading activity.

Of note, out of the 326 deletions and 385 insertions found in our MA-lines, only 1 insertion and 3 deletions were 2 bp in length, all others were of length 1. This differs from that reported for R$^{3610}$ (Sung et al. 2016) where (after mapping to our reference subset of positions in *B. subtilis* 168 for consistency with our data) 15 of the 64 deletions and 4 of the 24 insertions were of length >1, ranging from 2 to 30 (**Table S3**). We noted that the start or entirety of some of the longer deletions found in the R$^{3610}$ strains were identical to the start of the immediately following sequence (e.g., deletions at positions 491,610 and 959,732), suggesting that these deletions occurred by local recombination. To compare the similar events in the different strains, we discarded any indel of length > 2 in the R$^{3610}$ data from our calculation of indel rates. Also, an indel of size 2 was considered to result from a single deletion or insertion event.

#### 1.2 Genetic instability of inducible hypermutating synthetic circuits during MA experiments

Nonsynonymous mutations on the negative dominant *polC** or *mutL** alleles, or mutations on their RBS or promoter ($P_{hs}$), or nonsynonymous mutations on the *lacI* gene (which represses $P_{hs}$), as well as other mutations in untranslated regions of the *polC** and *mutL** mRNA that may alter their stability, will directly modify the mutation rate of *L**, *C**, and *LC** strains. Therefore, we established a list of the mutations identified on these genetic elements and in their vicinity. For lines $C^*_3$, $LC^*_2$, $LC^*_3$ and $LC^*_4$, we found nonsynonymous mutations on *polC** that could inactivate or decrease the activity of PolC*, allowing native PolC to replicate DNA better and correct mutations through its exonuclease activity. However, we did not find mutations in any of the above-mentioned genetic elements in the $LC^*_1$ line, and the appearance of the mutation found in $LC^*_3$ may be posterior to the decrease in mutation rate. This suggests the presence of compensatory mutations in the *B. subtilis* chromosome that can decrease substitution rates independently of the inducible synthetic hypermutation circuits.

In the strain *LC*,* all mutations on an inserted allele were found on *polC**, and none on *mutL**, consistent with the larger size of *polC** (4,311 bp) compared to *mutL** (1,884 bp). No mutation was found on the shorter genetic elements *lacI*, $P_{hs}$ or RBS of the mutant alleles.

#### 1.3 Disentangling the contributions of replication and transcription to mutation profiles

Due to the strong collinearity between transcription and replication in *B. subtilis*, apparent replication-oriented biases may involve a contribution from transcription. In fact and as expected, the asymmetry between template and non-template (also known as coding) strands of transcription is similar to the asymmetry between leading and lagging strands of replication (**Figure S11A**). To analyze whether the biases are related to replication, transcription, or both, we used a generalized linear model (GLM). This approach allowed us to disentangle the contributions of replication and transcription, and showed that once the replication strand was given, the transcription strand had no statistically significant effect on substitution rates (ANOVA *p*-value=0.11, considering all strains simultaneously). This is consistent with the comparison between estimates of local substitution rates distinguishing replication and transcription strands (**Figure 4B**) and also with previous statistical analyses on MMR-deficient strains (Schroeder et al. 2016). For example, regardless of the transcription strand, the substitution rate at C sites in the MMR-deficient strains is higher on the leading strand than on the lagging strand, and the substitution at T sites in the *LC** strain is higher on the lagging strand than on the leading strand. Further examination of these rates (**Figure 4B**) does not indicate a higher substitution rate in genes encoded "head-on" (*i.e.* where the template strand of transcription is the leading strand of replication), regardless of strain.

Regarding the location in or out of a coding region, a significant difference in mutation rates was found only for the $R^{3610}$ and $\Delta S^{3610}$ strains, with the $R^{3610}$ having a higher substitution rate in noncoding regions, whereas $\Delta S^{3610}$ has a higher substitution rate in coding regions (GLM, p-values $3.51\times10^{-9}$ and $3.53\times10^{-7}$, respectively, and **Figure S11B**) as also found in the analysis of (Sung et al., 2015). A higher G+C-content in coding than in noncoding regions combined with a globally higher substitution rate at sites corresponding to G:C pairs, and more generally a difference in nucleotide composition between coding and noncoding regions, may lead to a higher substitution rate in coding regions. In support of this idea, we found that a GLM including "triplet" and "coding status" does not fit the data for $\Delta S^{3610}$ significantly better than a GLM including only the "triplet" effect (ANOVA *p*-value>0.05). In contrast, a model including "triplet" and "coding status" fits the data for $R^{3610}$ better (ANOVA *p*-value=$1.04\times10^{-7}$).

The presence of the base in a coding or noncoding region could be refined using expression level data. However, the expression level of the region does not appear to affect the substitution rate in any strain (**Figure S11C**).

Taken together, these results lead us to conclude that the effect of transcription is absent or very marginal under our growth conditions.

#### 1.4 Bayesian estimation

All models were fitted using JAGS (Plummer, 2003) version 4.3.0 for posterior sampling via the R package rjags (https://cran.r-project.org/package=rjags). We used random initialization and discarded the first 4,000 iterations as burn-in. Convergence was assessed using the Gelman-Rubin statistic (Gelman and Rubin, 1992) with two independent Monte Carlo Markov chains. The total number of iterations was determined automatically using the runjags package (https://cran.r-project.org/package=runjags) which uses a minimum of 10,000 iterations and then extends the chain until the Gelman-Rubin statistic is less than 1.05 for all parameters. For all cases considered, 10,000 iterations appeared to be sufficient to achieve convergence. Each chain was thinned to a final size of 4,000 samples from the posterior distribution.

***Model without saturation***

The counts of mutations of the different types (or contexts) i observed in the MA-lines for strain s (“wild-type” R, “MMR-deficient” MMR-, “proofreading-deficient” *C**, “MMR-deficient and proofreading-deficient” *LC**) were modeled as independent Poisson distributed random variables with means n[i].μ[s,i].t[s], *i.e.*

m[s,i] ~ Poisson(n[i].μ[s,i].t[s]),

where n[i] is the number of possible sites for mutation of type i in the genome, μ[s,i] is the rate of mutation of type i in strain s per site per generation, and t[s] is the number of generations elapsed.

The mutation rates were modeled as independent log-normal random variables

μ[s,i] ~ log10 Normal(ν[i],σ[i]),

where ν[i] and σ[i] are strain-specific hyperparameters corresponding respectively to the location and scale parameters of the lognormal distribution (mean and standard deviation of the normal distribution). These hyperparameters are random variables drawn from the weakly informative hyperpriors

ν[i] ~N(mean=−8,standard deviation=3),

σ[i]/3 ~ Student+(degrees of freedom=5).

This choice of a truncated heavy-tailed Student's t-distribution, suggested in (Gelman, 2006), allow shrinkage when compatible with the data, and is intended to be more robust than the classical inverse gamma prior.

#### 1.5 Mathematical modeling of MMR saturation

Mechanistically, the rate of mutation of type i per site per generation in the four strains was modeled as

μ[LC*,i]=γ[i],

μ[MMR-,i]=γ[i]$q_{Proofreading}$[i],

μ[C*,i]=γ[i](θ+(1-θ).$q_{MMR}$[i]),

μ[wt,i]=γ[i]$q_{Proofreading}$[i]$q_{MMR}$[i],

where θ is a mixture parameter common to all values of i, γ[i] is the error rate before correction by proofreading or MMR, $q_{Proofreading}$[i] is the rate at which errors of type i escape proofreading correction, and $q_{MMR}$[i] is the rate at which errors of type i escape MMR correction.

***Bayesian estimation of the model with MMR saturation in the strain C****

A uniform prior on the interval [0,1] was used for θ.

For μ[LC*,i], μ[MMR-,i], and μ[wt*,i] the model was exactly the same lognormal model as described above in the context of the model without saturation of the MMR while μ[C*,i] was written

μ[C*,i]=μ[LC*,i](θ+(1-θ).μ[wt,i]/μ[MMR-,i]).

***Proportion of the observed mutations in the strain C* that were not subjected to MMR correction***

The parameter θ corresponds to the proportion of errors that occurred in a context or saturated MMR in strain *C**. Notably, since these errors are never corrected by the MMR they account for a proportion of the observed mutations in this strain greater than θ. This proportion, represented in the inset plot of **Figure 7A**, is given by

$\theta/(\theta+(1-\theta).q_{MMR}[i])$,

where $q_{MMR}[i]=\mu[wt,i]/\mu[MMR-,i]$.

***Relationship between θ and the maximum number of errors that the MMR can handle***

If we assume that λ corresponds to the maximum number of errors that can be handled by the MMR in one replication of the bacterial chromosome and that the number of errors submitted to correction by the MMR follows a Poisson distribution with mean κ, then the proportion of errors that are not subject to MMR correction because they occur after saturation writes as a function of κ and λ

$\eta(\kappa,\lambda)=(\kappa.q_\kappa(\lambda-1)-\lambda q_\kappa(\lambda))/\kappa$,

where $q_\kappa$ denotes the tail function of the Poisson distribution with mean κ, *i.e.* $q_\kappa(l)=1-\Sigma_{n\leq l}\exp(-\kappa)\kappa^n/n!$.

To understand the origin of this relationship, note that the mean of the Poisson distribution truncated to n>λ writes $\kappa.q_\kappa(\lambda-1)/q_\kappa(\lambda)$, where κ is the mean of the untruncated distribution. In a draw of this truncated distribution, the average number of errors that are not subject to MMR correction because they occur after saturation is $\kappa.q_\kappa(\lambda-1)/q_\kappa(\lambda)-\lambda$. This quantity multiplied by $q_\kappa(\lambda)$, the probability of having a draw greater than λ, and divided by κ, the total number of errors, gives the proportion η(κ,λ).

In strain *C**, $\kappa=\Sigma_i\gamma[i].n[i]$ and thus θ, which corresponds to the proportion of errors that occurred in a context or saturated MMR, writes $\theta=\eta(\Sigma_i\gamma[i].n[i],\lambda)$. This relationship can be numerically inverted to obtain λ as a function of θ and $\Sigma_i\gamma[i].n[i]$ (as represented in **Figure 7B**).

In the wild-type, κ is equal to $\Sigma_i\gamma[i].n[i].q_{Proofreading}[i]$ and to $\Sigma_i\mu[MMR-,i].n[i]$. With the mutation rates observed in the experimental data, the corresponding η(κ,λ) is very close to 0.

***Relationship between θ and the distance the polymerase must travel after a first error before MMR can handle a second error.***

An alternative mechanism can lead to MMR saturation: the requirement of a minimum distance between two errors corrected by the MMR. This minimum distance is hereafter denoted by R in analogy with the refractory period of a neuron. Assuming that polymerase errors occur as a Poisson process with rate γ, the errors subject to MMR correction follow a renewal process with inter-event distance R+1/γ. The average ratio between the number of events subject to MMR correction and the total number of errors, which corresponds to 1-θ, is then 1/(1+γR).

Thus, we can derive R from μ and θ using the inverse relationship $R=(1/\gamma).[\theta/(1-\theta)]$.

Considering $\gamma=7.4\times10^{-7}$ $bp^{-1}$ (the total point mutation rate in *LC**) and the point estimates of θ between 0.071 and 0.084 (depending on the data set, estimated in *C**), we obtain a value of R between 103 and 124 kbp.

#### 1.6 Algebraic analysis of a general model with two repair pathways and two subclasses of errors

We consider here a model with two subclasses of errors, numbered 1 and 2, and in which the wild-type possess two error correction pathways, denoted a and b. The mutation rates are denoted μ[a,b] for the wild-type, μ[-,b], μ[a,-], and μ[-,-], for pathways a and b deactivated separately or simultaneously. In this model the mutation rates write

μ[-,-]=γ[1]+γ[2],

μ[a,-]=γ[1].q[a,1]+γ[2].q[a,2],

μ[-,b]=γ[1].q[b,1]+γ[2].q[b,2],

μ[a,b]=γ[1].q[a,1].q[b,1]+γ[2].q[a,2].q[b,2],

where γ[1]≥0 and γ[2]≥0 correspond to the initial rates of both subclasses of errors, q[a,1]∈(0,1) and q[a,2]∈(0,1) are the rates at which the errors of subclass 1 and 2, respectively, escape correction by pathway a, and q[b,1]∈(0,1) and q[b,2]∈(0,1) are the rates at which the errors of subclass 1 and 2, respectively, escape correction by pathway b.

This model has 6 free parameters and can perfectly fitany set of four mutation rates that satisfy the four “natural” order relations μ[-,-]≥μ[a,-], μ[-,-]≥μ[-,b], μ[-,a]≥μ[a,b], and μ[-,b]≥μ[a,b], and the condition that the increments in the mutation rates are at least additive which can be written

μ[-,-]≥μ[a,-]+μ[-,b]-μ[a,b] or

μ[-,-]≥(μ[a,-]-μ[a,b])+(μ[-,b]-μ[a,b])+μ[a,b].

In practice, to find a set of parameters that fits a particular set of mutation rates we can for example, choose q[a,1]=0 and q[a,2]=1 (pathway ‘a’ corrects perfectly the errors of subclass 1 and none of subclass 2) and then identify

γ[1]=μ[-,-]-μ[a,-]

γ[2]=μ[a,-]

q[b,1]=(μ[-,b]-μ[a,b])/(μ[-,-]-μ[a,-])

q[b,2]=μ[a,b]/μ[a,-]

***Special cases***

In the case with a single error subclass (γ[1]=0 or γ[2]=0), we have

μ[-,-]/μ[a,b]=(μ[a,-]/μ[a,b]).(μ[-,b]/μ[a,b]).

We can see in this equation that the fold-changes by which the mutation rates increase with the deactivation of each correction pathway are multiplicative.

The same property of multiplicative fold-changes holds in another special case where a pathway does not distinguish between the two subclasses of errors (q[a,1]=q[a,2] or q[b,1]=q[b,2]).

On the contrary, if each pathway corrects only a specific subclass of errors (e.g. q[a,2]=1 and q[b,1]=1, meaning that ‘a’ cannot correct subclass 2 and ‘b’ cannot correct subclass 1), then

μ[-,-]-μ[a,b]=(μ[-,b]-μ[a,b])+(μ[a,-]-μ[a,b]).

In this case the increments in the mutation rate associated with the deactivation of each correction pathway are additive.

***Extreme scenarios of super-multiplicative and sub-multiplicative ratios***

The condition for multiplicative fold-changes is

μ[-,-]/μ[a,b]=(μ[a,-]/μ[a,b]).(μ[-,b]/μ[a,b]),

and can be rewritten

R=(μ[-,-].μ[a,b])/(μ[a,-].μ[-,b])=1,

where R>1 corresponds to “super-multiplicative” fold-changes and R<1 corresponds to “sub-multiplicative” fold-changes.

To explore algebraically how to obtain extreme values of R, this ratio can be written in terms of the 6 model parameters

R=(γ[1]$^2$q[a,1]q[b,1]+γ[1]γ[2]q[a,1]q[b,1]+γ[1]γ[2]q[a,2]q[b,2]+γ[2]$^2$q[a,2]q[b,2])/(γ[1]$^2$q[a,1]q[b,1]+γ[1]γ[2]q[a,1]q[b,2]+γ[1]γ[2]q[a,2]q[b,1]+γ[2]$^2$q[a,2]q[b,2]).

We note that R does not change when γ[1] and γ[2] are multiplied by the same constant, or q[a,1] and q[a,2] are multiplied by the same constant, or q[b,1] and q[b,2] are multiplied by the same constant. Without loss of generality, to find the extreme values of R, we can then set γ[1]=γ and γ[2]=1-γ, q[a,1]=α and q[a,2]=1-α, and q[b,1]=β and q[b,2]=1-β. The ratio R can then be rewritten as a function of γ, α, and β as

R(γ,α,β)=[γ.α.β+(1-γ).(1-α).(1-β)]/[(γ.α+(1-γ).(1-α)).(γ.β+(1-γ).(1-β))].

We can then do a logarithmic transformation (monotone) of the function R(γ,α,β) and study the first order partial derivatives ∂log(R)/∂α, and ∂log(R)/∂β. The partial derivative ∂log(R)/∂α is null if and only if γ(1-γ)(2β-1)=0 and has the sign of 2β-1. Symmetrically, the partial derivative ∂log(R)/∂β is null if and only if γ(1-γ)(2α-1)=0 and has the sign of 2α-1.

We deduce from these derivatives that the ratio is constant over the values of (α,β) when γ=0 or γ=1, which corresponds to a scenario with a single non-empty subclass. We have already pointed out that the property of multiplicative fold-changes holds in this special case (R(0,α,β)=1 and R(1,α,β)=0).

We also see from these derivatives that for any given value of γ, the ratio is constant over the values of α if β=1/2 and reciprocally over the values of β if α=1/2. We have already pointed out that the property of multiplicative fold-changes holds in this special case (R(γ,α,1/2)=1 and R(γ,1/2,β)=1).

By examining the sign of the two first order partial derivatives ∂log(R)/∂α and ∂log(R)/∂β we see that for any given value of γ such as 0<γ<1, the ratio increases when (α,β)->(1,1) (or symmetrically (α,β)->(0,0)) and decreases when (α,β)->(1,0) (or symmetrically (α,β)->(0,1)). The corresponding extrema can be computed and are $R(\gamma,0,1)=0$ and $R(\gamma,1,1)=1/\gamma$ (symmetrically $R(\gamma,1,0)=0$ and $R(\gamma,1,1)=1/(1-\gamma)$).

The minimum value of R is 0 and is reached in a special case of additive increments where the two repair systems are both complementary and perfectly efficient $\mu[a,b]=0$.

The maximum value of R for $0<\gamma<1$ is $1/\gamma$ if $\gamma<1/2$ and $1/(1-\gamma)$ if $\gamma>1/2$. In this scenario, the two repair pathways correct the same subclass of errors and R can be increased indefinitely by decreasing the size ($\gamma$ or $(1-\gamma)$) of the subclass of errors that are not corrected by either of the two considered pathways. In this way, the model can account for arbitrary super-multiplicative fold-changes.

#### 2 Supplementary Figures

##### 2.1 Figure S1

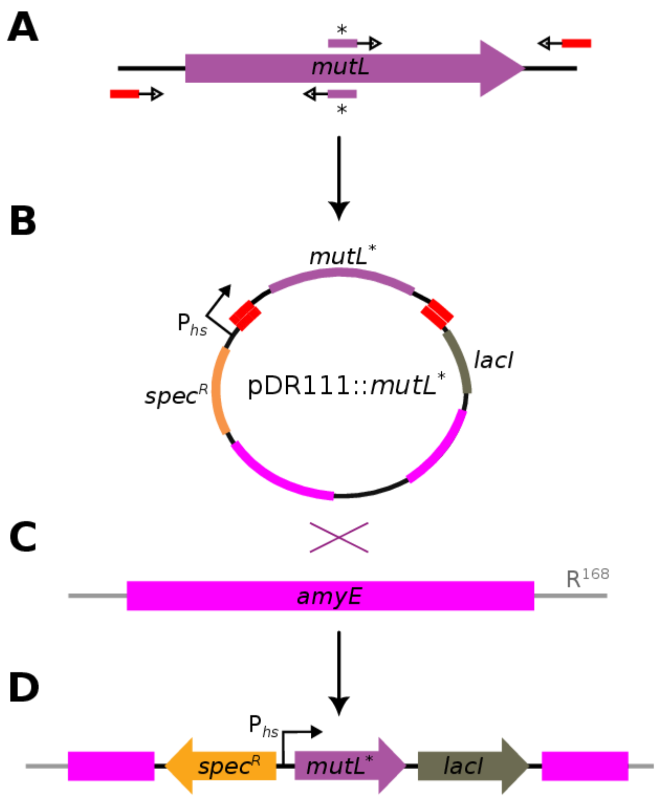

**Figure S1. Steps for the introduction of a mutant allele into the chromosome of *B. subtilis* using pDR111. A.** PCR amplification to introduce the desired point mutation N34H into *mutL*. **B.** Insertion into pDR111. **C.** Insertion into the *B. subtilis* genome by homologous recombination in *amyE.* **D.** Final insert in the chromosome of the strain *L**. A similar procedure was used to obtain *C** and *LC**, but with *polC mut1* amplified directly from a strain carrying this mutant allele of *polC*.

#### 2.2 Figure S2

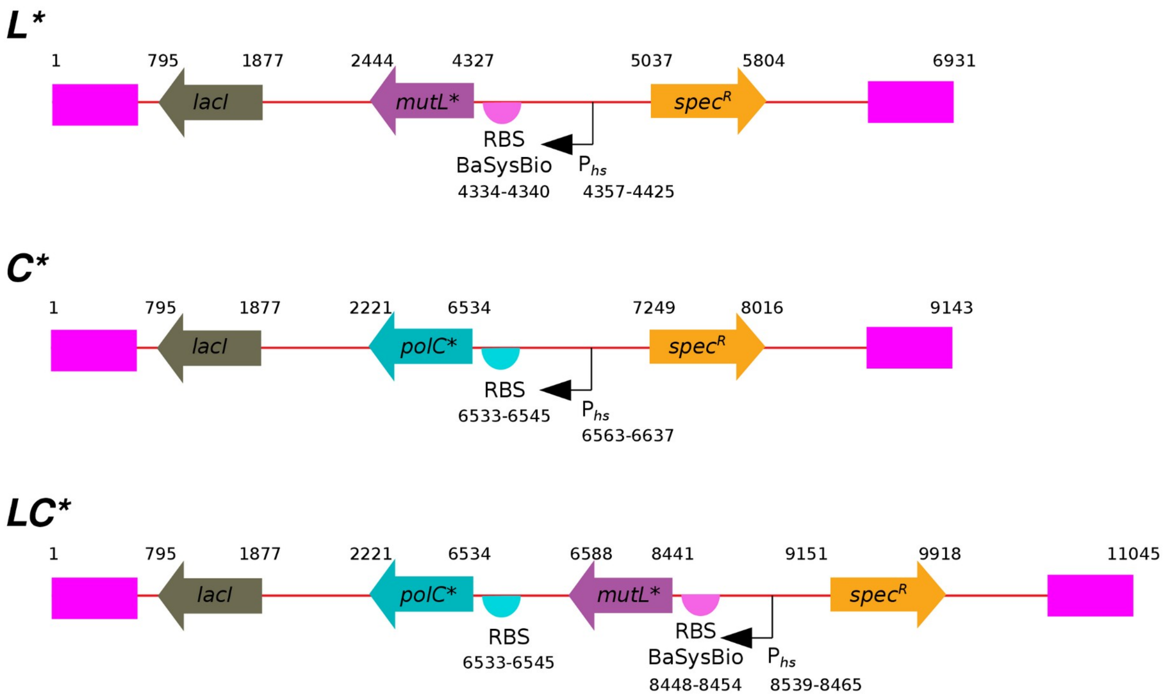

**Figure S2. Schematic representation of the inserts carrying the hypermutator inducible circuits.** Regions of *amyE* are represented in magenta. Coordinates of the different genes, promoters and RBSs are given in the three strain-specific inserts.

#### 2.3 Figure S3

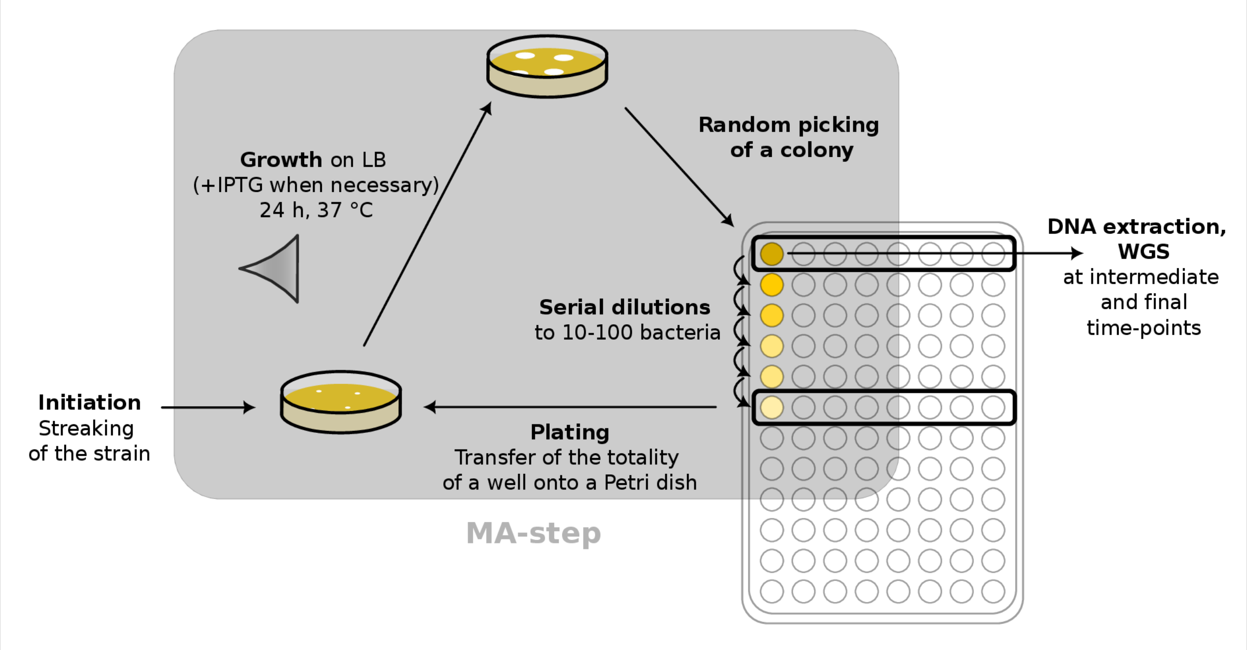

**Figure S3. Mutation accumulation protocol.** The use of a 96-well plaque allows parallel serial dilution of independent MA-lines. Each well in the first row of the 96-well plaque contains all the cells of a single colony; the mutations identified by sequencing are those that occurred before the clonal expansion of that colony (i.e. before the current MA-step).

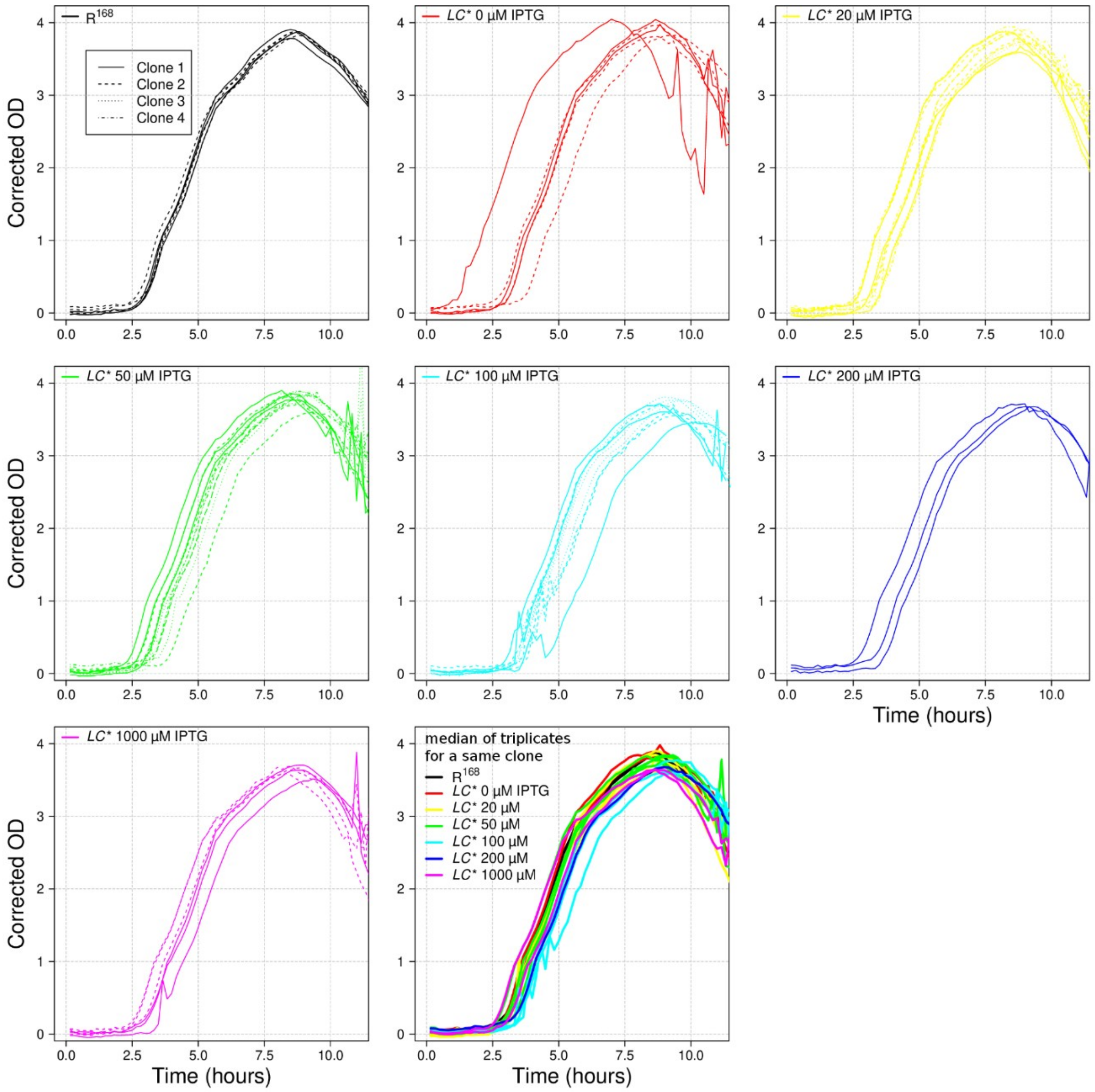

#### 2.4 Figure S4

**Figure S4. Growth rates of strain *LC** at increasing IPTG concentrations and comparison with $R^{168}$.** Corrected optical density (microplate reader measured OD) at 600 nm during growth in LB at 37°C. Triplicate growth curves for several isolated clones (1 to 4) are shown for different combinations of strains and conditions in the first seven plots. In the last plot, the combinations of strains and conditions are all represented together, with each line corresponding to the median of the growth curves for the same clone.

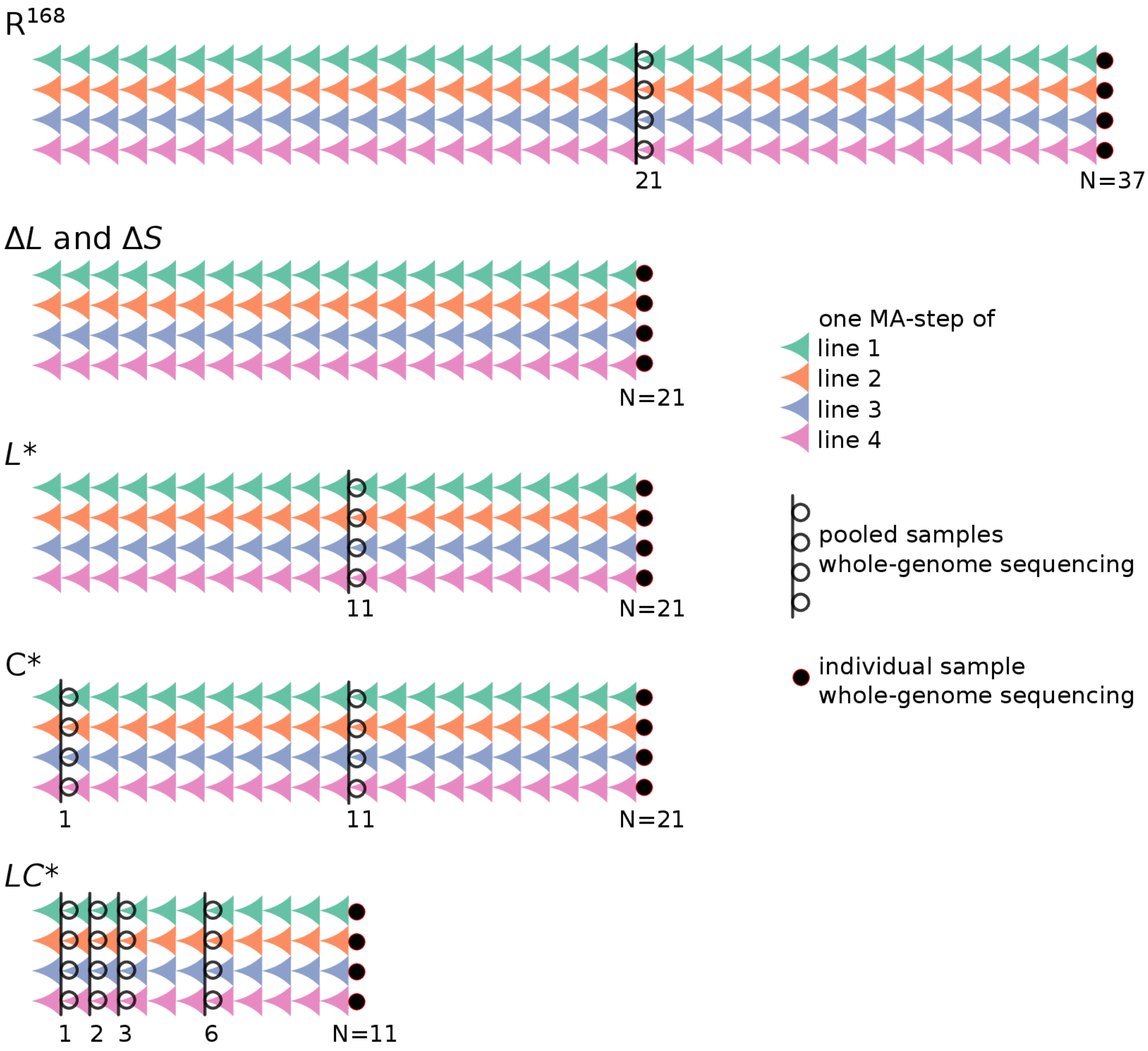

#### 2.5 Figure S5

**Figure S5. Design of mutation accumulation experiments.** For each strain the four independent MA-lines are represented, with MA-steps symbolized by concave triangles (clonal expansion during colony growth). The total number of MA-steps (N) and the time points of whole-genome sequencing are reported. Sequencing data from pooled samples identify the time interval in which a mutation occurred.

#### 2.6 Figure S6

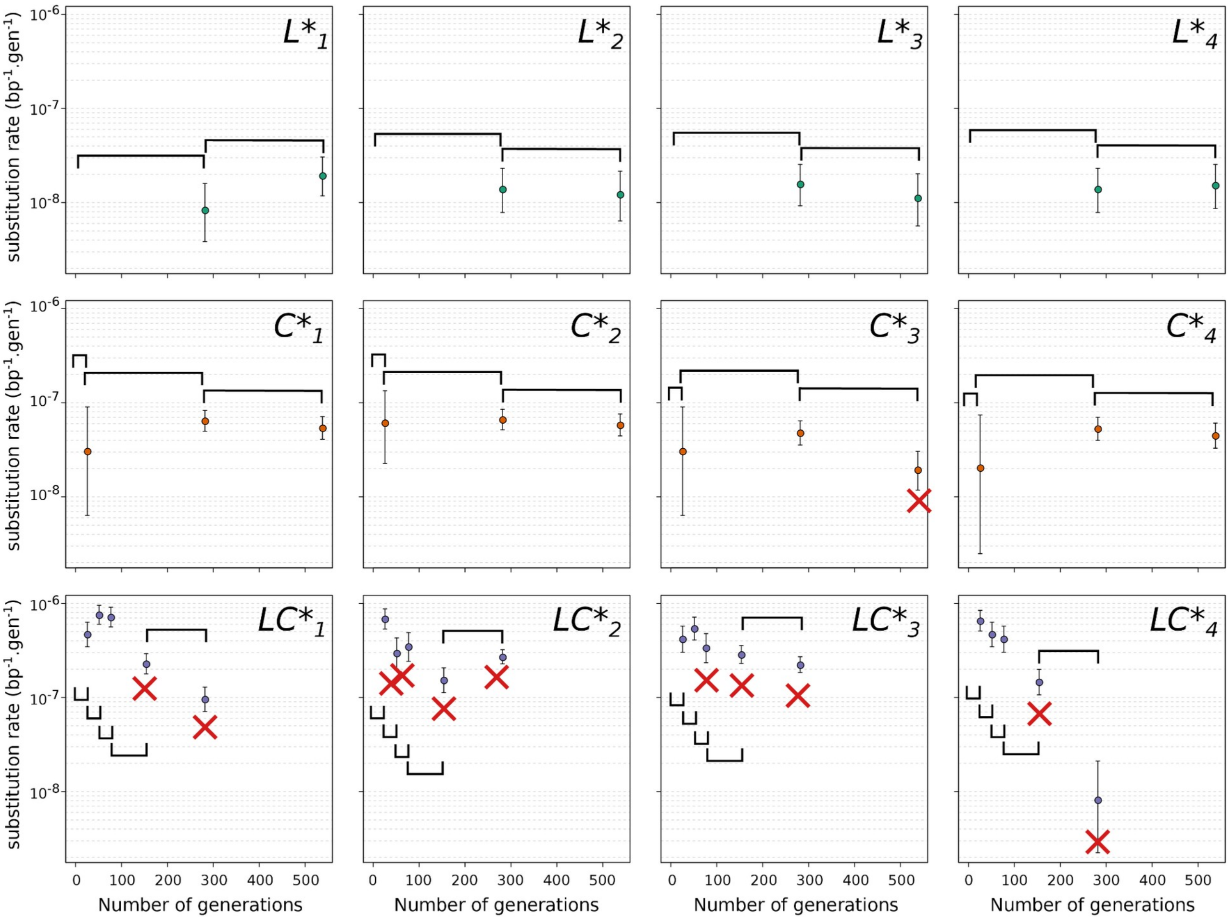

**Figure S6. Evolution of substitution rates during mutation accumulation experiments.** Substitution rates per base and per generation for the four MA lines of the *L** (top), *C** (middle) and *LC** (bottom) strains are calculated using the number of indels fixed within each interval. 95% confidence intervals are represented. The time intervals discarded due to a decrease in substitution rate are indicated with red crosses.

#### 2.7 Figure S7

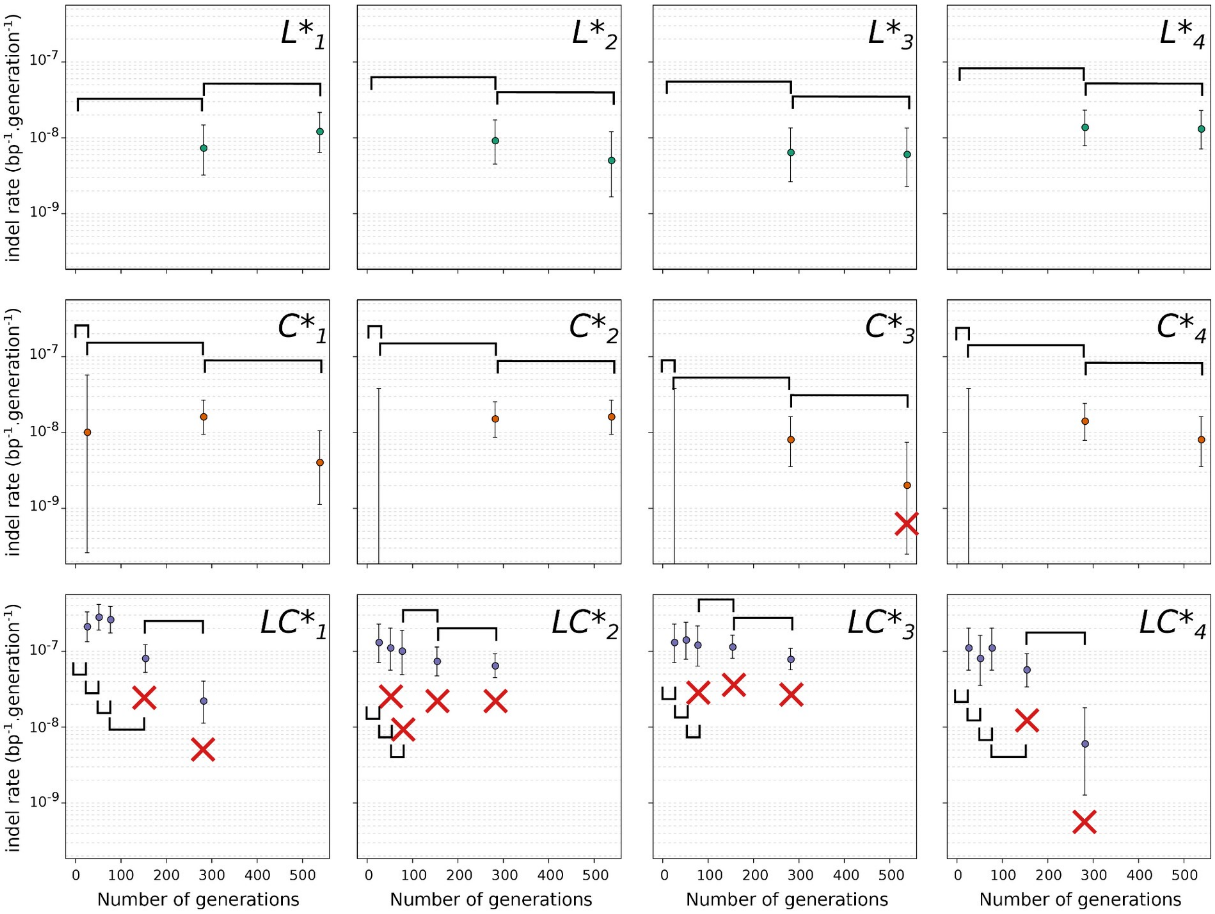

**Figure S7. Evolution of indel rates in Mutation Accumulation experiments.** Indel rates per base and per generation for the four MA lines of the *L** (top), *C** (middle) and *LC** (bottom) strains are computed using the number of indels fixed within each interval. 95% confidence intervals are represented. The time intervals discarded due to a decrease in substitution rate are indicated with red crosses.

#### 2.8 Figure S8

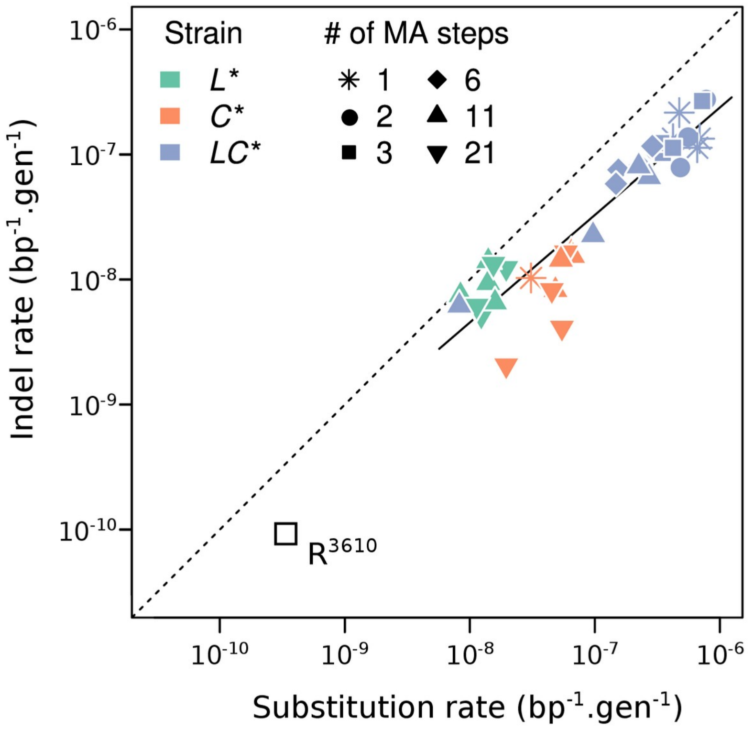

**Figure S8. Indel versus substitution rates for each MA-line and interval.** The black line represents the fit of a linear model of log(indel rates) vs. log(substitution rates). The Pearson correlation coefficient is 0.93 (p-value=$2.28\times10^{-16}$). The point representing the indel and substitution rates of $R^{3610}$ is added for comparison.

#### 2.9 Figure S9

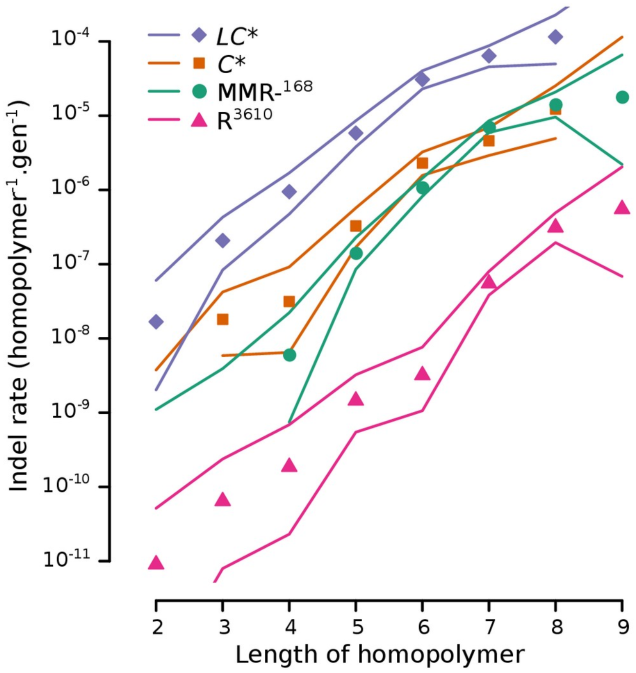

**Figure S9. Indel rates as a function of homopolymer length.** The indel rate in a homopolymer of a given length is given per homopolymer occurrence and per generation.

#### 2.10 Figure S10

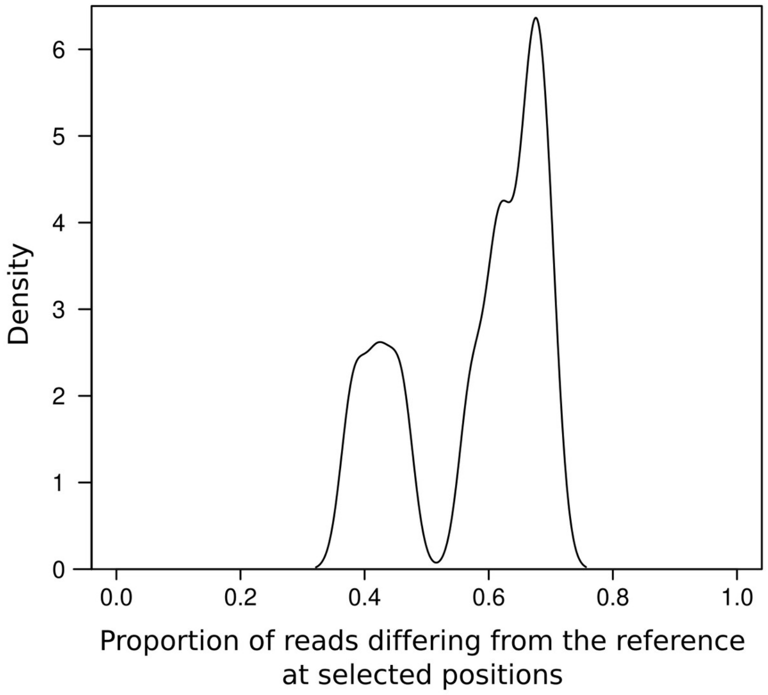

**Figure S10. Bimodal distribution of the proportion of reads differing from the reference for reads mapped in the inducible synthetic circuits.** Density estimation based on substitutions at the 10 positions in *mutL*, *mutL**, *polC* or *polC** listed in Table S3 with a Gaussian kernel smoother (standard deviation = 0.2). The bimodal distribution reflects a higher gene dosage for *mutL** and *polC** than for *mutL* and *polC* because the *amyE* locus where the mutant alleles are inserted is closer to the replication origin of the chromosome. Mutations associated with an alternative allele frequency above 50% are in *mutL** and *polC** as verified by PCR for some of the mutations (see **Table S4**).

#### 2.11 Figure S11

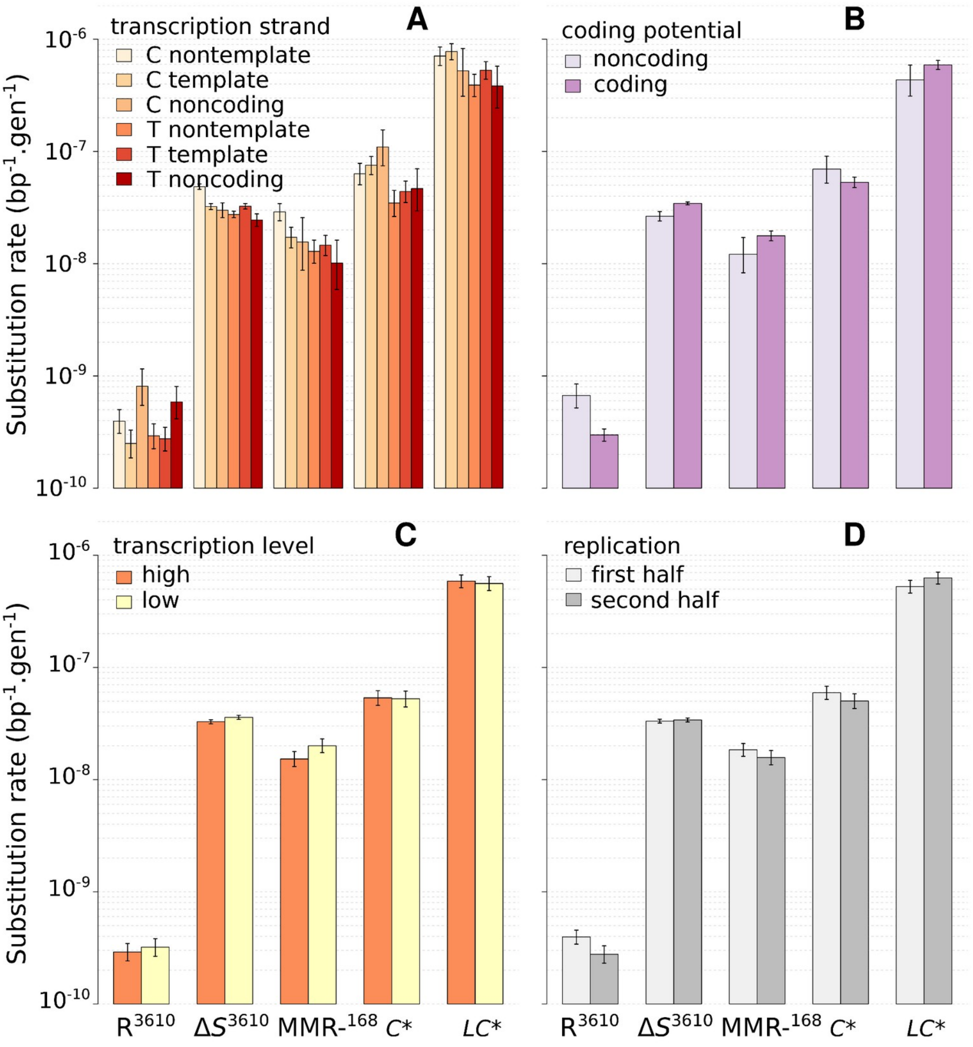

**Figure S11. Substitution rates in different chromosomal contexts.** Error bars represent the 95% confidence interval according to the Poisson distribution. **A.** Effect orientation with respect to the transcription strand, the pyrimidine of the pair determines the strand of a mutation site. **B.** Effect of localization in coding or noncoding regions. **C.** Effect of transcription level. **D.** Effect of distance from the origin of replication.

#### 2.12 Figure S12

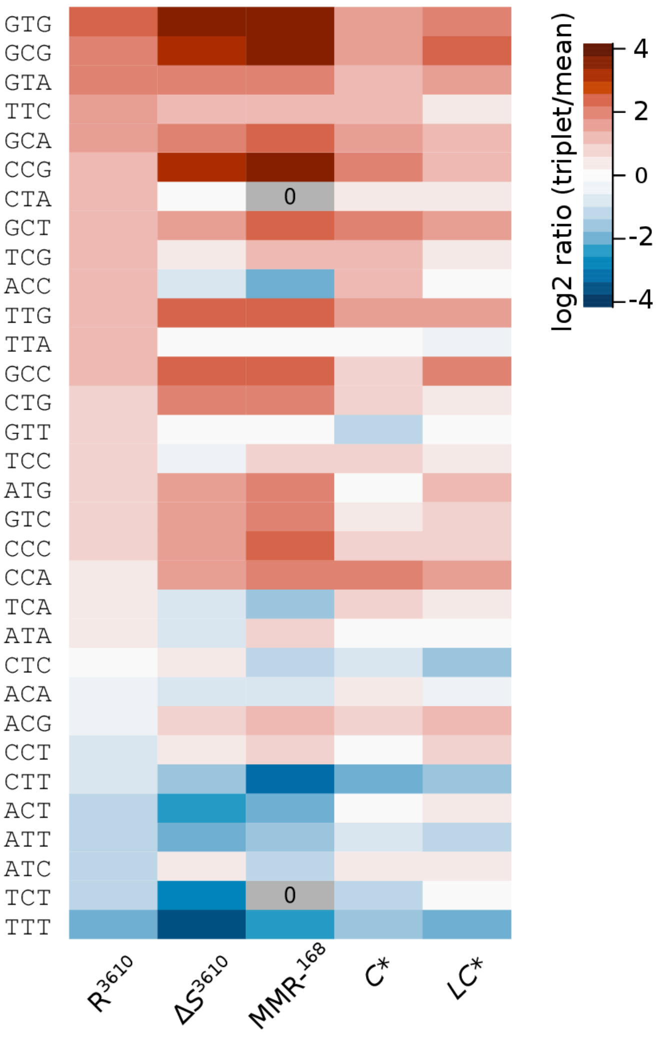

**Figure S12. Heatmap representation of the heterogeneity of substitution rates among triplets.** Each triplet corresponds to the mutated pyrimidine and its 5' and 3' nucleotides. The log2 ratio of the estimated rate for the triplet wrt to the mean for each genotype is represented. Rates were estimated by maximum likelihood, absence of substitutions in a triplet context is indicated by a “0” in a gray cell. Triplets are ordered vertically according to the substitution rate in the wild-type ($R^{3610}$, leftmost column).

#### 2.13 Figure S13

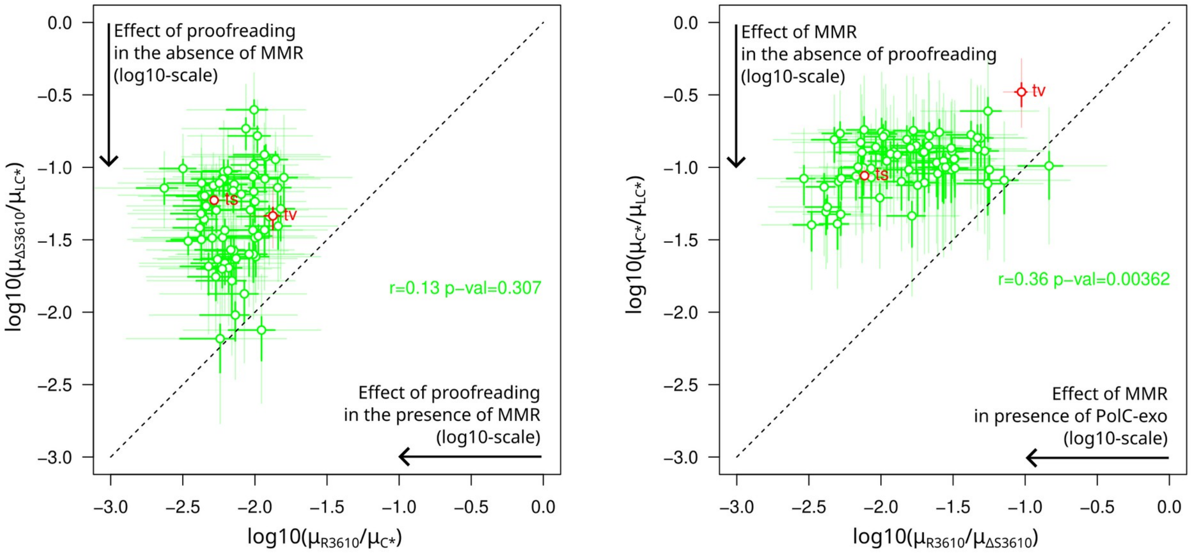

**Figure S13. Effects of proofreading and MMR in the presence or absence of the other system, based on** ***ΔS*****[3610].** Left plot: effect of proofreading in the presence or absence of MMR. Right plot: effect of activating MMR in the presence or absence of proofreading. This figure is similar to main text **Figure 6** but the MMR-deficient substitution profile is represented here by Δ*S*[3610] instead of MMR-[168].

#### 2.14 Figure S14

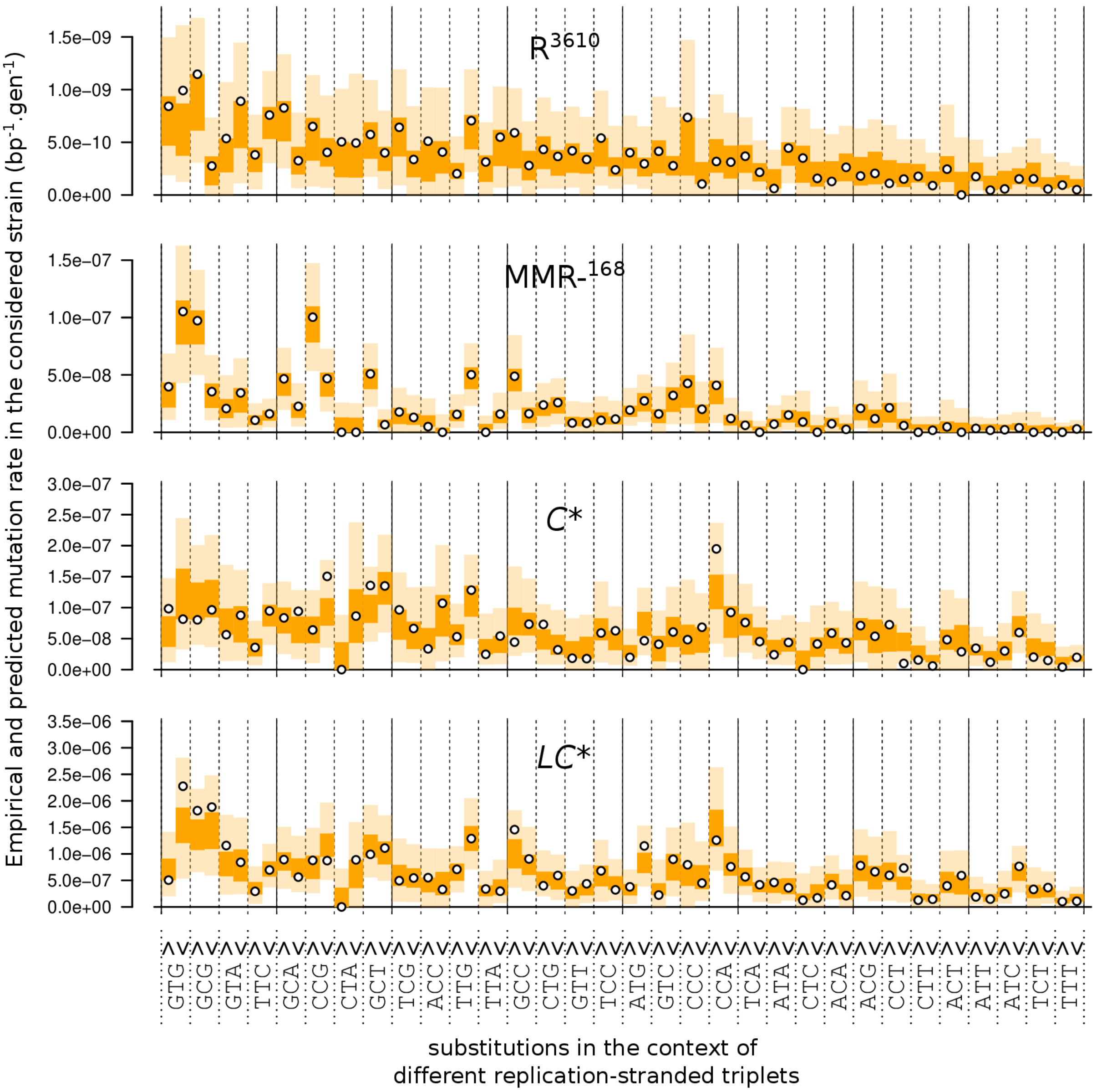

**Figure S14. Assessment of the fit of the MMR-saturation model to the experimental data considering the substitution profile obtained for MMR-$^{168}$.** Points represent empirically calculated substitution rates, *i.e.* the number of observed substitutions divided by the number of possible sites in the genome and the number of generations. Colored areas represent the distribution of the empirical rates simulated under the posterior distribution of the model parameters (50% of the density in the darker areas, 95% if the lighter areas are also considered). Replication-stranded triplets are ordered in decreasing order of non-stranded empirical substitution rates and then pyrimidine on the leading and lagging strands of replication.

#### 2.15 Figure S15

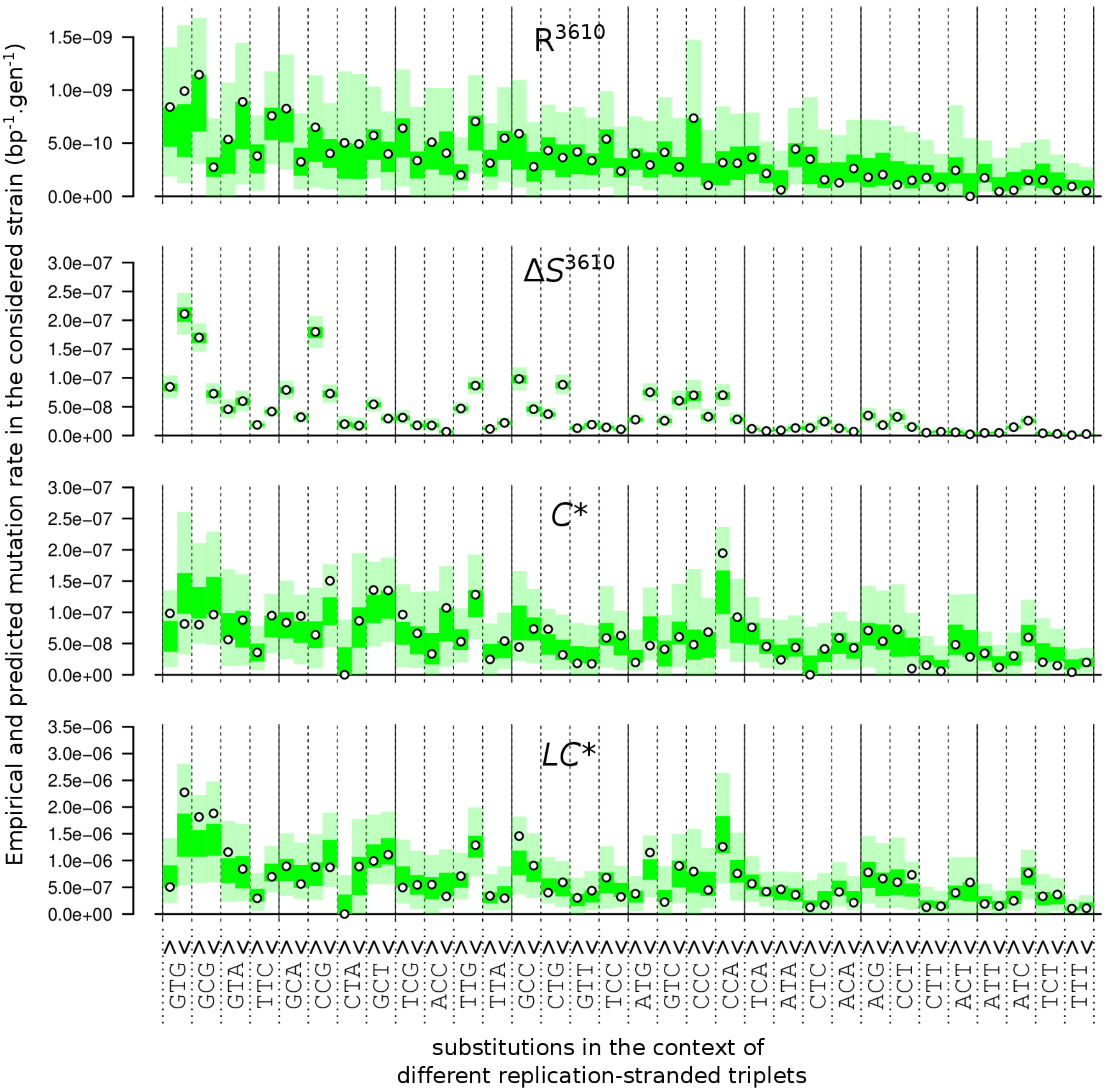

**Figure S15. Assessment of the fit of the MMR-saturation model to the experimental data considering the substitution profile obtained for $\Delta S^{3610}$.** Same as **Figure S14** but the data used for MMR-deficient substitution profile is here $\Delta S^{3610}$ instead of MMR-$^{168}$.

#### 2.16 Figure S16

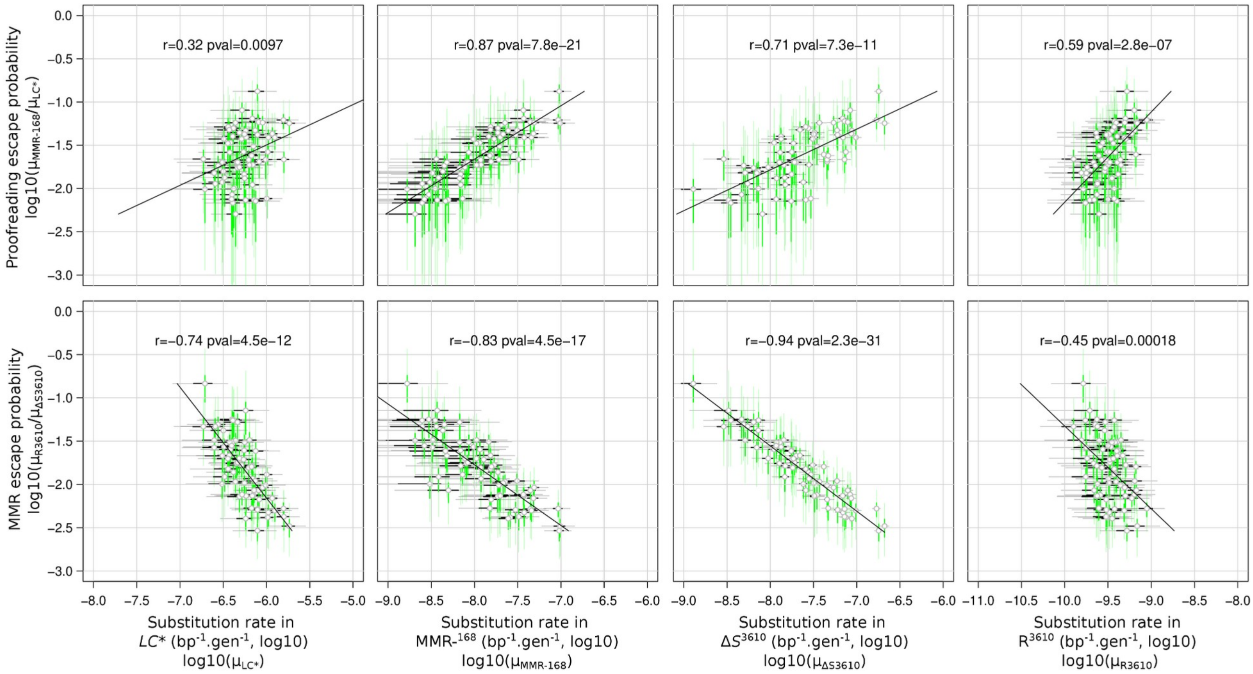

**Figure S16. Correlations between proofreading or MMR escape probability and substitution rates in different strains.** Each point represents a replication-oriented triplet. Around each point, the 50% and 95% marginal credibility intervals on the horizontal and vertical axes, computed from the quantiles of the posterior distributions, are represented by segments (bold and dark vs. thin and light, respectively). Pearson correlation coefficients computed between escape probability and substitution rate (both on logarithmic scale) are reported with the corresponding p-value. The best-fit line obtained by linear regression is also shown.

#### 2.17 Figure S17

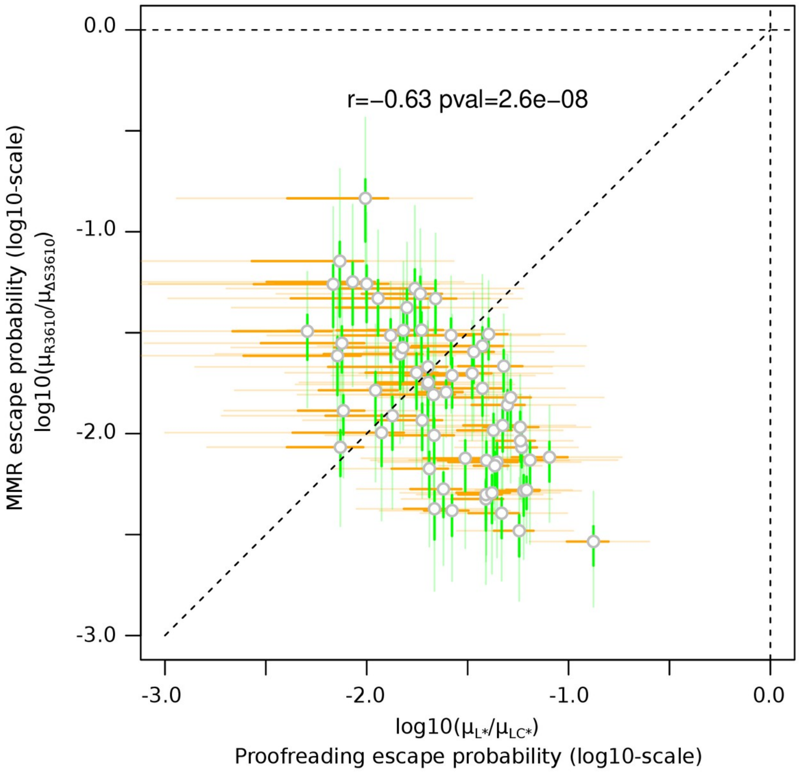

**Figure S17. Comparison of the apparent efficiency of MMR and proofreading correction of errors in replication-stranded triplets.** The efficiency of correction by MMR shown here was estimated from the data for background *B. subtilis* 3610 collected by Sung *et al.* (2016). The proofreading activity considered here is the activity abolished in the *polC** allele, and was obtained by comparing the substitution profiles of strains *L** and *LC** (IPTG 100 µM). Despite the uncertainty of individual measures for the 64 different stranded-triplets (bold and thin bars represent 50% and 95% credibility intervals along each axis), a strong overall negative correlation is detected (Pearson correlation coefficient r=-0.63, *p*-value=$2.6\times10^{-8}$).

### 3 Supplementary Tables

#### 3.1 Table S1

##### Table S1. Detailed results of fluctuation assays.

| Strain | [IPTG] (µM) | #cultures[a] | #zeros[b] | mean M[c] | V[d] (mL) | mean C[e] ($\times10^7$) | sd C[f] ($\times10^7$) | MLE m[g] | RifR mutation rate[h] |
|---|---|---|---|---|---|---|---|---|---|
| $R^{168}$ | 0 | 96 | 84 | 0.14 | 1.0 | 12.95 | 6.66 | 0.13 | $9.74\times10^{-10}$ |
| $R^{168}$ | 100 | 96 | 76 | 0.25 | 0.2 | 7.99 | 7.60 | 0.21 | $2.68\times10^{-9}$ |
| *ΔS* | 0 | 16 | 0 | 37.75 | 1.0 | 12.95 | 6.66 | 9.64 | $7.54\times10^{-8}$ |
| *ΔL* | 0 | 16 | 0 | 50.69 | 1.0 | | | 11.79 | $9.10\times10^{-8}$ |
| *L** | 0 | 8 | 6 | 0.25 | 1.0 | 11.82 | 5.25 | 0.25 | $2.12\times10^{-9}$ |
| | 5 | 9 | 7 | 0.22 | 0.2 | 4.48 | 2.25 | 0.22 | $4.85\times10^{-9}$ |
| | 20 | 8 | 0 | 36.63 | 1.0 | 11.82 | 5.25 | 11.21 | $9.48\times10^{-8}$ |
| | 50 | 9 | 0 | 27.00 | 0.2 | 3.60 | 2.50 | 8.14 | $2.26\times10^{-7}$ |
| | 100 | 8 | 0 | 506.38 | 1.0 | 11.82 | 5.25 | 43.19 | $3.66\times10^{-7}$ |
| | 500 | 8 | 0 | 191.38 | 1.0 | | | 40.88 | $3.46\times10^{-7}$ |
| *C** | 0 | 9 | 6 | 0.33 | 0.2 | 4.58 | 2.25 | 0.33 | $7.27\times10^{-9}$ |
| | 5 | 9 | 7 | 1.67 | 0.2 | | | 0.27 | $5.81\times10^{-9}$ |
| | 20 | 9 | 0 | 8.67 | 0.2 | | | 1.76 | $3.83\times10^{-8}$ |
| | 50 | 9 | 0 | 60.56 | 0.2 | | | 16.41 | $3.58\times10^{-7}$ |
| | 100 | 9 | 0 | 228.22 | 0.2 | | | 45.73 | $9.98\times10^{-7}$ |
| | 500 | 9 | 0 | 459.67 | 0.2 | | | 82.65 | $1.80\times10^{-6}$ |
| *LC** | 0 | 9 | 6 | 0.56 | 0.2 | 3.60 | 2.50 | 0.39 | $1.09\times10^{-8}$ |
| | 5 | 9 | 5 | 4.56 | 0.2 | | | 0.55 | $1.52\times10^{-8}$ |
| | 20 | 9 | 0 | 32.44 | 0.2 | | | 9.99 | $2.77\times10^{-8}$ |
| | 50 | 9 | 0 | 1166.00 | 0.2 | | | 178.09 | $4.94\times10^{-6}$ |
| | 100 | 9 | 0 | 1404.89 | 0.2 | | | 208.13 | $5.78\times10^{-6}$ |
| | 500 | 9 | 0 | 1351.56 | 0.2 | | | 202.25 | $5.61\times10^{-6}$ |

[a] Number of cultures for RifR determination. [b] Number of samples without any RifR mutants. [c] Mean number of mutants per culture. [d] Culture volume (mL), the dilution factor used to estimate the total number of cells was $20\times10^5$ for 1 mL cultures $2\times10^5$ for 0.2 mL cultures. [e-f] Mean and standard deviation of the final number of cells (calculated by day) ($\times10^7$). [g] MLE of the number of mutations per culture. [h] Rate of emergence of the RifR phenotype estimated from the number of mutations and the number of cells per culture.

#### 3.2 Table S2

##### Table S2. Induction of *mutL** and *polC** by IPTG as measured by RNA-Seq.

| Str[a] | [IPTG] (μM) | R[b] | *mutL+L** (fpkm)[c] | mutL* N34H[d] | *polC+C** (fpkm)[c] | *polC** G430E[d] | *polC** S621N[d] |
|---|---|---|---|---|---|---|---|
| R[168] | 0 | 1 | 104.3 | 0.00 (0.00-0.02) | 159.5 | 0.00 (0.00-0.02) | 0.00 (0.00-0.03) |
| R[168] | 0 | 2 | 106.1 | 0.00 (0.00-0.03) | 159.5 | 0.00 (0.00-0.01) | 0.00 (0.00-0.02) |
| R[168] | 100 | 1 | 96.5 | 0.00 (0.00-0.02) | 144.5 | 0.00 (0.00-0.02) | 0.00 (0.00-0.03) |
| R[168] | 100 | 2 | 116.7 | 0.00 (0.00-0.02) | 171.5 | 0.00 (0.00-0.01) | 0.00 (0.00-0.02) |
| *L** | 100 | 1 | 7,020.1 | 0.99 (0.99-1.00) | 134.4 | 0.00 (0.00-0.05) | 0.00 (0.00-0.07) |
| *L** | 100 | 2 | 7,234.1 | 0.99 (0.98-0.99) | 108.2 | 0.00 (0.00-0.03) | 0.00 (0.00-0.04) |
| *C** | 100 | 1 | 72.3 | 0.00 (0.00-0.04) | 4,471.8 | 0.98 (0.97-0.98) | 0.97 (0.97-0.98) |
| *C** | 100 | 2 | 74.5 | 0.00 (0.00-0.05) | 3,905.8 | 0.97 (0.97-0.98) | 0.98 (0.98-0.99) |
| *LC** | 0 | 1 | 148.8 | 0.35 (0.28-0.42) | 197.0 | 0.18 (0.13-0.23) | 0.25 (0.18-0.32) |
| *LC** | 0 | 2 | 98.6 | 0.29 (0.22-0.38) | 123.7 | 0.19 (0.13-0.27) | 0.28 (0.19-0.40) |
| *LC** | 100 | 1 | 3,601.8 | 0.98 (0.98-0.99) | 3,084.8 | 0.97 (0.97-0.98) | 0.96 (0.95-0.97) |
| *LC** | 100 | 2 | 5,905.3 | 0.98 (0.98-0.98) | 5,356.4 | 0.96 (0.96-0.97) | 0.97 (0.96-0.97) |

[a] Tested strain. [b] RNA-Seq biological replicate. [c] Quantification of the amount of transcript for this gene (wild-type and mutant allele) in fpkm (fragments per kilobase of transcript per million mapped reads). [d] Proportion of RNA-Seq reads carrying the mutation out of the reads overlapping this position.

#### 3.3 Table S4

**Table S4. Aggregated numbers of indels, indel rate and proportion of insertions for each investigated strain.**

| Strain | Indels[a] | | Indel rate [95% CI] | Proportion of insertions [95% CI] |
|---|---|---|---|---|
| | insertions | deletions | | |
| $R^{168}$ | 0 | 3 | $4.2 \times 10^{-10}$ [$0.86\text{-}12 \times 10^{-10}$] | 0.00 [0.00-0.71] |
| $R^{3610}$ | 20 | 53 | $9.2 \times 10^{-11}$ [$7.2\text{-}12 \times 10^{-11}$] | 0.27 [0.18-0.39] |
| $\Delta S^{3610}$ | - | - | - | - |
| $\Delta L$ | 24 | 58 | $10 \times 10^{-9}$ [$8.0\text{-}12 \times 10^{-9}$] | 0.29 [0.20-0.40] |
| $\Delta S$ | 31 | 37 | $8.3 \times 10^{-9}$ [$6.5\text{-}11 \times 10^{-9}$] | 0.46 [0.33-0.58] |
| *L** | 23 | 53 | $9.3 \times 10^{-9}$ [$7.3\text{-}12 \times 10^{-9}$] | 0.30 [0.20-0.42] |
| MMR- | 78 | 148 | $9.2 \times 10^{-9}$ [$8.1\text{-}11 \times 10^{-9}$] | 0.35 [0.28-0.41] |
| *C** | 47 (2) | 35 (0) | $1.1 \times 10^{-8}$ [$0.91\text{-}1.4 \times 10^{-8}$] | 0.57 [0.46-0.68] |
| *LC** | 95 (125) | 50 (90) | $1.7 \times 10^{-7}$ [$1.4\text{-}2.0 \times 10^{-7}$] | 0.66 [0.57-0.73] |

[a] Between parentheses: number of indels in time intervals with decreased mutation rates (discarded from the analysis).

#### 3.4 Table S5

**Table S5. Mutations in MA-lines found in sequencing reads that mapped on the inserted regions for *L**, *C** and *LC** strains.**

| MA line | MA-step[a] | Position[b] | Region[c] | Ref. | Alt[d] | Gene | AA | Consequence |
|---|---|---|---|---|---|---|---|---|
| *L** 1 | 11 | 3244 | native | C | -1T | *mutL* | 361+ | frameshift |
| *C** 3 | 11 | 4683 | insert | C | T | *polC* | 618 | nonsynonymous (GCC→ACC) |
| *C** 3 | 11 | 4477 | insert | G | A | *polC* | 686 | synonymous (TTC→TTT) |
| *C** 4 | 11 | 6296 | insert | G | A | *polC* | 80 | nonsynonymous (TCT→TTT) |
| *C** 4 | 11 | 8014 | insert | T | -1A | *spec*[R] | 255+ | frameshift |
| *LC** 1 | 11 | 404 | insert | C | T | intergenic | - | - |
| *LC** 1 | 11 | 696 | insert | G | A | intergenic | - | - |
| *LC** 1 | 1 | 8592 | insert | T | +1A | intergenic | - | - |
| *LC** 2 | 3 | 5325 | insert$ | C | T | *polC* | 404 | nonsynonymous (GGC→AGC) |
| *LC** 2 | 6 | 6217 | insert$ | C | T | *polC* | 106 | synonymous (CAG→CAA) |
| *LC** 2 | 6 | 2455 | native$ | G | A | *polC* | 1360 | synonymous (TCC→TCT) |
| *LC** 2 | 6 | 1932 | insert | T | G | intergenic | - | - |
| *LC** 3 | 6 | 4251 | insert$ | T | C | *polC* | 762 | nonsynonymous (AAT→GAT) |
| *LC** 3 | 11 | 5345 | native$ | T | C | *polC* | 397 | nonsynonymous (GAA→GGA) |
| *LC** 4 | 6 | 6399 | insert | A | G | *polC* | 46 | nonsynonymous (TGG→CGG) |
| *LC** 4 | 3 | 4245 | native | G | A | *polC* | 764 | nonsynonymous (CAT→TAT) |

[a] MA-step after which the sequencing that allowed the first detection was performed. [b] Position of the mutation as seen when the sequencing read is mapped to the insert (see **Figure S2**) [c] Mutations detected in *mutL* and *polC* could have occurred either in the native allele or in the mutant allele located in the insert. For these two genes, the information in this column is derived from the frequency of the mutation in the reads (see **Figure S7,** experimentally verified by PCR amplification and sequencing when indicated by the symbol $). [d] Observed substitution, deletion (-) or insertion (+).

#### 3.5 Table S7

##### Table S7. Primers used for the construction of strains.

| Name | Sequence [a] | Role |
|---|---|---|
| P1 | 5′–GCTCGTTGCACACACCATTT–3′ | Amplification of *mutSL* operon – F |
| P2 | 5′–TGAAAAAGGCCCTTCCCCAG–3′ | Amplification of *mutSL* operon – R |
| P3 | 5′–ACTAAGCTTAATTGTTATCCGCTCA–3′ | Linearization pDR111 – F |
| P4 | 5′–GCATGCAAGCTAATTCGGTGG–3′ | Linearization pDR111 - R |
| P5 | 5′–GGATAACAATTAAGCTTAGTCGGGAAGGAGGAACTACTGTGGCAAAAGTCATCCAACTGT–3′ | Amplification of *mutL* for Gibson assembly in pDR111 – F |
| P6 | 5′–CCGAATTAGCTTGCATGCTCACACGAACAGGGAGCAAA–3′ | Amplification of *mutL* for Gibson assembly in pDR111 -R |
| P7 | 5′–*CGTCAAAGAATTGGTGGAA**C**ATGCGATCGACGCTGACAGCA*CAGTCATTG–3′ | Introduction of N34H mutation on *mutL* – F |
| P8 | 5′–*TGCTGTCAGCGTCGATCGCAT**G**TTCCACCAATTCTTTGACG*ACTGAGGCGG –3′ | Introduction of N34H mutation on *mutL* – R |
| P9 | 5′–GGATAACAATTAAGCTTAGTTTTAGGGAGGGATACTGTCT–3′ | Amplification of *polC** for Gibson assembly in pDR111 – F |
| P10 | 5′–CCGAATTAGCTTGCATGCCCTAAGCAAGTGACAGAAACT–3′ | Amplification of *polC** for Gibson assembly in pDR111 – R |
| P11 | 5′–CTACATCACGCGTTTGAAC–3′ | Amplification of *mutL** for Gibson assembly with *polC** - R |
| P12 | 5′–TGTTCAAACGCGTGATGTAGTTTTAGGGAGGGATACTGTCT–3′ | Amplification of *polC** for Gibson assembly with mutL* - F |

[a] Regions used for the HiFi DNA assembly protocol are underlined. The single nucleotide mutations used to build the *mutL*(N34H) variant is indicated in bold. The sequences in primers P7 and P8 that allow for the assembly by PCR are indicated in italics.
